# Hatcheries to high seas: climate change connections to salmon marine survival

**DOI:** 10.1101/2023.09.18.558187

**Authors:** Shuichi Kitada, Katherine W. Myers, Hirohisa Kishino

## Abstract

We investigated variations in the marine survival of Japanese hatchery chum salmon (*Oncorhynchus keta*) during 25 years of climate change (1998–2023). Japan is the world’s largest producer of hatchery salmon, and is located near the global southern distribution limit of chum salmon. Our goal was to identify local and context-specific metrics related to the observed coastwide decline in salmon marine survival over the past two decades. We hypothesized multiple metrics in three categories of stressors: hatchery carryovers, ocean conditions, and predators and competitors. The hatchery carryovers are stressors related to hatchery rearing that affect survival at a different life stage. We collected, processed, and collated large publicly available datasets into a comprehensive open-access database encompassing the life cycle of Japanese chum salmon, from eggs to adult spawners. Multivariate regression models showed associations between stressors and adult salmon return rate (marine survival) varied by coastal management region, salmon life stage, and seasonal high-seas distribution area. In the early marine life history stage, parental egg size and fry size-at-release had the largest positive model effects on marine survival. The sea surface temperature (SST) at time of fry release and a predator of fry had significant negative effects. In the offshore and high-seas life stages, summer SST had negative effects, while winter SST had positive effects. Russian chum and/or pink salmon abundance had negative effects, while no effect was found for North American pink and chum salmon abundance. Generalized additive models (GAMs) identified a nationwide decline in egg size and fry size-at-release. Our study highlights the need for an experimental approach to hatchery practices, including monitoring and analyses with updated information, leading to effective management decisions and policies for future sustainability and conservation of salmon resources.

## 1| INTRODUCTION

Climate change influences the survival and abundance of many economically important marine and anadromous fish species, including Pacific salmon (*Oncorhynchus* spp.) (Springer & van Vliet, 2014; Cheung & Frölicher, 2020; Cheung et al., 2021; Crozier et al., 2021; Crozier & Siegel, 2023). Pacific salmon populations often consist of hundreds of subpopulations with diverse life history traits and local adaptations to environmental variation (Groot & Margolis, 1991; Hilborn et al., 2003; Larson et al., 2014; Quinn, 2018). Pacific salmon species are distributed in coastal marine waters and throughout the high seas (international waters) of the North Pacific Ocean and Bering Sea, where Asian and North American populations intermingle during seasonal feeding migrations (Myers et al., 2007; Beamish, 2018; Quinn, 2018). Their wide geographic range, long ocean migrations, and complex life histories expose salmon populations to multiple climate-driven stressors at both local and North Pacific-wide scales (Cline et al., 2019; Crozier et al., 2021). Global warming of 1.5°C above pre-industrial levels is projected to shift the ranges of many marine species to higher latitudes and increase damage to marine ecosystems, such as rising temperatures, ocean acidification, heatwaves, decreases in sea ice, habitat loss, disruptions in food webs, and declining fish populations (Dahlke et al., 2018; Frölicher et al., 2018; Masson-Delmotte, 2022).

The effects of climate change, ocean acidification, and large-scale hatchery releases on the abundance and productivity of Pacific salmon populations have received increasing attention in recent years (Lewis et al., 2015; Jeffrey et al., 2017; Cunningham et al., 2018; Cline et al., 2019; Losee et al., 2019; Connors et al., 2020, 2024; Oke et al., 2020; Ohlberger et al., 2022, 2023; Malick et al., 2023). Declines in body size, primarily due to a shift in age structure towards younger ages at maturity, were found in chum, Chinook (*O. tshawytscha*), coho (*O. kisutch*), and sockeye (*O. nerka*) salmon from large-scale analyses across Alaska and were linked to climate change and increased inter- and intra-specific competition due to an overabundance of salmon in the high seas (Lewis et al., 2015; Oke et al., 2020; Ohlberger et al., 2023). Pink salmon (*O. gorbuscha*) abundance increased since the mid-1970s to record highs in the 2000s, supported by climate change effects on the ocean environment and production processes, and may have affected the abundance of salmon, other species, and their ecosystems (Springer & van Vliet, 2014; Connors et al., 2020; Ruggerone et al., 2023).

Pacific salmon have a long history of artificial propagation beginning in the mid-1800s, and their management comprises the largest set of hatchery release programs (Naish et al., 2007). Chum salmon hatcheries in Japan constitute the world’s largest salmon hatchery release program (Amoroso et al., 2017; Kitada, 2018), continuing for over 130 years (Miyakoshi et al., 2013). Hatchery releases of fry chum in Japan increased rapidly in the 1970s, but gradually declined from a peak of 2 billion per year in 1983 to 1.3 billion per year in 2021 because of poor returns of adult spawners. Adult returns of Japanese hatchery chum increased rapidly since the 1970s, but declined sharply after the late 1990s from the historical maximum of 88 million in 1996 to the minimum of 19 million in 2021. Previous studies have reported higher spring SSTs in the coastal zone contributed to better survival of juvenile chum through improved feeding, growth, and swimming ability (Saito & Nagasawa, 2009; Kuroda et al., 2020; Honda et al., 2021). Conversely, shorter inshore stays associated with warming SSTs could lead to lower juvenile survival and insufficient growth (Kaeriyama, 2022; Honda et al., 2025). Although it is generally accepted that a warming climate is affecting marine survival of salmon, the mechanisms underlying recent declines in Japanese hatchery chum salmon are largely unknown. The need for adaptive management is advocated for the long-term sustainability of chum salmon (Kaeriyama, 2008, 2022; Kaeriyama et al., 2012).

The development of adaptive management plans to address climate-change impacts on fish, fisheries, aquaculture, and associated habitats and ecosystems requires knowledge of “local and context-specific indicators associated with climate stressors” (FAO, 2022). Adaptive management is the design of management actions as experiments with feedback through monitoring, enabling an iterative process of learning and adapting to uncertainties and risks (Walters & Hilborn, 1978; Hilborn & Sibert, 1988; Bahri et al., 2021; Bryndum-Buchholz et al., 2021). A comprehensive review of literature addressed strengths and weaknesses of climate-change research by life stage and showed the complexity of whole life cycle approaches to adaptive management strategies for anadromous salmonids (Crozier & Siegel, 2023). Here, we investigated associations between adult return rates (an index of marine survival) of Japanese hatchery chum salmon by life stage and a suite of hypothesized climate-change stressors. In our study, we expected survival of all life stages of chum salmon to be associated with climate change impacts on coastal and open ocean ecosystems. Our primary objective was to identify potential stressors along known coastal and high-seas migration routes of Japanese chum salmon for use in regional adaptive management of salmon resources.

## 2| MATERIALS AND METHODS

### 2.1 Ecological setting, study design, and methods overview

The ecological setting of our study encompasses the known geographic distribution and seasonal migration patterns of Japanese chum salmon in the North Pacific Ocean and adjacent seas (Figure 1; reviewed by Fisheries Agency of Japan, 1955; Sano, 1966, 1967; Shepard et al., 1968; Neave et al., 1976; Salo, 1991; Urawa et al., 2018). Chum salmon fry are released from more than 260 coastal hatcheries in northern Honshu and Hokkaido in winter and spring (Kitada, 2020, Figure 1a) and migrate rapidly through river estuaries (Kobayashi, 1980; Kaeriyama, 1989; Saito & Nagasawa, 2009). After ocean entry, fry migrate northward in shallow coastal waters before moving offshore to summer feeding grounds in the southern Sea of Okhotsk (Figure 1b; Fukuwaka & Suzuki, 1998; Urawa et al., 2006; Kasugai et al., 2016; Shubin & Akinicheva, 2016; Honda et al., 2025). Japanese chum salmon spend their first ocean winter (age-1) in the western North Pacific Ocean and in spring move northeastward to summer feeding grounds in the Bering Sea (Figure 1b; Mayama & Ishida, 2003; Urawa et al., 2018). At subsequent life stages (ages 2–4; immature and maturing), they make seasonal migrations between the Gulf of Alaska in winter and the Bering Sea in summer (Figure 1b; Beacham et al., 2009; Urawa et al., 2009, 2016). In spring, the northward migration of maturing chum salmon precedes that of immature fish (Neave et al., 1976; Nagasawa & Azumaya, 2009). In late summer and fall, maturing fish return to Japan from the Bering Sea through the western North Pacific (Figure 1b; Yonemori, 1975; Urawa et al., 2018).

**FIGURE 1.**
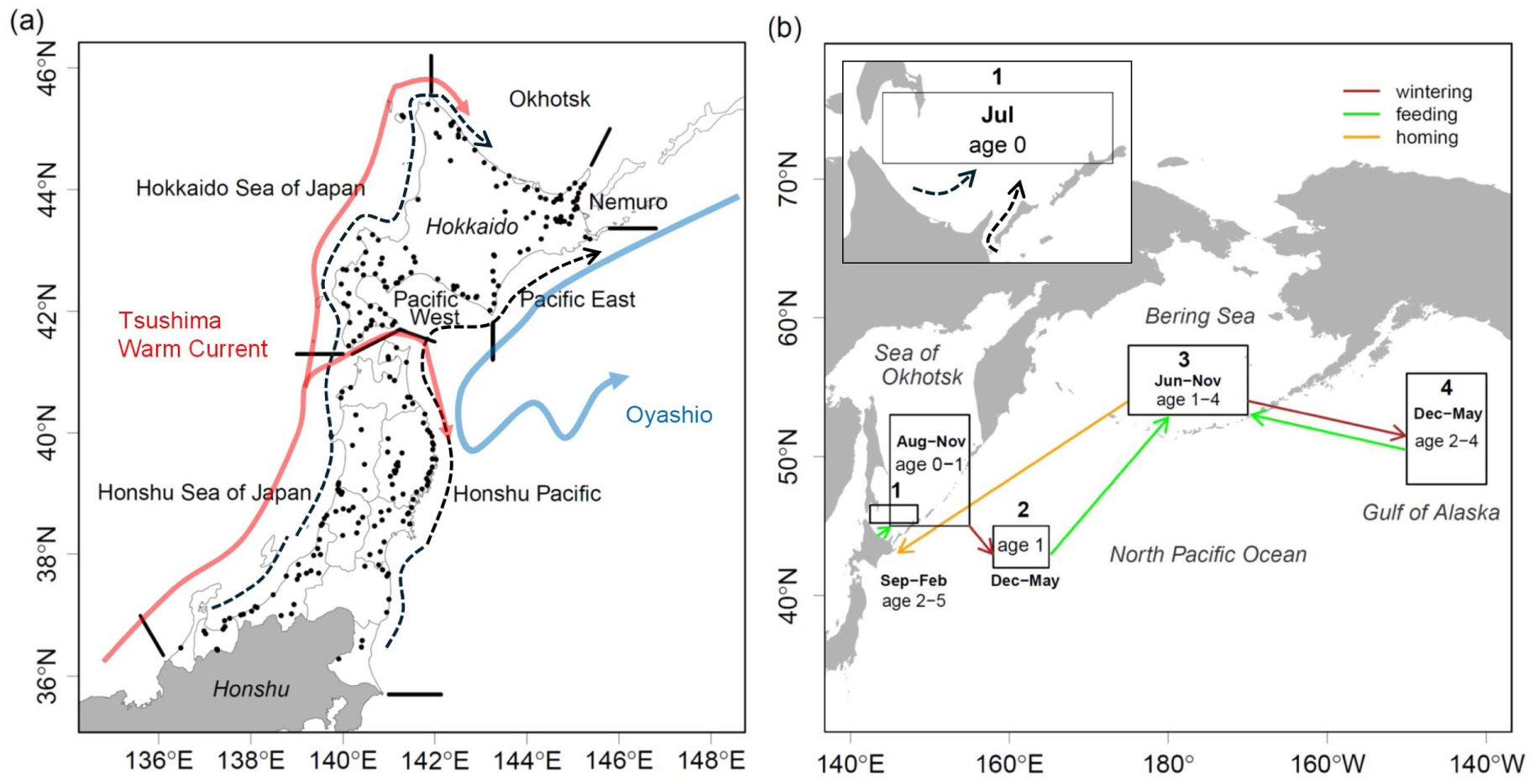
Japanese chum salmon management regions, coastal migration routes, and high-seas distribution areas. (a) Seven management regions (delineated by black bars) and hatcheries (black dots) in Japan. Migration paths of hatchery-released fry were modified from Saito and Nagasawa (2009, Figure 1). Schematic sea surface currents were redrawn from Kuroda et al. (2020, Figure 1). (b) principal areas of high-seas distribution (boxes), month(s), age group(s), and seasonal direction of wintering, feeding and homing migrations (arrows), numbered areas are: 1.Southern Sea of Okhotsk (45.2°N–46.5°N, 142.5°E–148.5°E), 2. Western North Pacific (42°N–45°N, 158°E–165°E), 3. Central Bering Sea (53°N–58°N, 175°E–170°W), 4. Central Gulf of Alaska (48°N–56°N, 140°W–150°W). Migration routes of Japanese chum salmon estimated from long-term field surveys, modified from Myers et al. (2007, Figure 1) and Urawa et al. (2018, Figure 37).

A brief overview of our study design and methodology for evaluating relationships between salmon marine survival and hypothesized stressors is provided below (see details in sections 2.2–2.8). Our study design is observational (a retrospective case study). We identified multiple *a priori* hypotheses about potential stressors driving marine survival of Japanese hatchery chum salmon at specific life stages and locations along their coastal and high-seas migration routes (Table 1). We grouped hypothesized stressors into three broad categories: hatchery carryovers, ocean conditions (SST), and predators and competitors. Hatchery carryovers are stressors related to hatchery rearing that affect survival at a different life stage. We assembled available data sets for use as measures of hypothesized stressors in each category. We summarized time series data on reproductive and biological traits of hatchery salmon and adult return rates by coastal management region (Figure 1a). We created isotherm maps to illustrate decadal-scale trends in SST throughout the known ocean range of Japanese chum salmon. We estimated SSTs at ocean entry and along coastal and high-seas migratory routes (Figure 1b). We developed measures of potential coastal marine fish predators and high-seas salmon competitors of Japanese chum salmon. Investigation of coastal mammal and bird predators and high-seas predators was beyond the scope of our analysis due to inadequate time-series data covering the period of our study. We estimated annual adult return rates (marine survival), stratified by salmon age group and year. We used multivariate regression models to evaluate relationships between the marine survival of salmon and hypothesized categories of stressors by coastal management region in Japan and seasonal high-seas habitats in the Sea of Okhotsk, western North Pacific, central Bering Sea, and Gulf of Alaska. Readers are cautioned that our statistical modeling results are correlational, not causal.

**TABLE 1.** Three subset models of stressors related to chum salmon marine survival (adult return rate) of fry released in year *t* (1998–2019) and explanatory variables considered in a regression model and hypothesized directional relationship. Sea surface temperature (SST); Western North Pacific (WN Pacific); Gulf of Alaska (GOA).

| Subset model | Explanatory variable (year) | Age<sup>†</sup> | Spatial scale<sup>‡</sup> | Hypothesized relationship to chum salmon survival (source) |
| --- | --- | --- | --- | --- |
| Hatchery carryovers | Parental relative fecundity ($t-1$) | 0 | Japan coast | Survival related (negative) to higher fecundity (Hasegawa et al., 2021a) |
| | Egg size ($t$) | 0 | Japan coast | Survival related (positive) to larger eggs (Hasegawa et al., 2021a) |
| | Fry size at release ($t$) | 0 | Japan coast | Survival related (positive) to size at release (Hasegawa et al., 2021b) |
| Ocean conditions | SST at release ($t$) | 0 | Japan coast | Survival related (negative) to SST (Nagata et al., 2007) |
| | Okhotsk coast Jul SST ($t$) | 0 | Okhotsk | 1<sup>st</sup> summer survival related (negative) to SST (Nagata et al., 2007) |
| | WN Pacific Jan-Apr SST ($t+1$) | 1 | WN Pacific | 1<sup>st</sup> winter survival related (positive) to SST (this study) |
| | Bering Sea Aug-Sep SST ($t+1$) | 1 | Bering Sea | 2<sup>nd</sup> summer survival related (negative) to SST (this study) |
| | GOA Jan-Apr SST ($t+2$) | 2 | Gulf of Alaska | 2<sup>nd</sup> winter survival related (positive) to SST (this study). |
| Predators and competitors | Yellowtail catch ($t$) | 0 | Japan coast | 1<sup>st</sup> year survival related (negative) to predation (Beamish, 2018) |
| | Russia chum catch ($t+3$) | 0–4 | Okhotsk, North Pacific | 1<sup>st</sup> - 3<sup>rd</sup> year survival related (negative) to Russian chum (this study). |
| | Russia pink catch ($t+1$) | 0–1 | Okhotsk, WN Pacific | 1<sup>st</sup> year survival related (negative) to Russian pink (this study) |
| | Russia pink catch ($t+2$) | 1–2 | Bering Sea | 2<sup>nd</sup> year survival related (negative) to Russian pink (Ruggerone et al., 2023) |
| | Russia pink catch ($t+3$) | 2–3 | Gulf of Alaska, Bering Sea | 3<sup>rd</sup> year survival related (negative) to Russian pink (Ruggerone et al., 2023) |
| | US & Canada chum catch ($t+3$) | 1–4 | North Pacific | 2<sup>nd</sup> - 3<sup>rd</sup> year survival related (negative) to US & Canada chum (this study) |
| | US & Canada pink catch ($t+1$) | 0–1 | Okhotsk, WN Pacific | 1<sup>st</sup> year survival related (negative) to US & Canada pink (this study) |
| | US & Canada pink catch ($t+2$) | 1–2 | Bering Sea | 2<sup>nd</sup> year survival related (negative) to US & Canada pink (this study) |
| | US and Canada pink catch ($t+3$) | 2–3 | Gulf of Alaska, Bering Sea | 3<sup>rd</sup> year survival related (negative) to US & Canada pink (this study) |

<sup>†</sup> Age of Japanese chum salmon released in year $t$ (=1998,...,2019). <sup>‡</sup> See Japanese salmon migration route (Figure 1).

### 2.2 Data sources, sample periods, sample sizes

We assembled available time-series datasets during 25 years of climate change (1998–2023) from public open-data websites. We assembled longer data time series to aid in interpreting results.

We obtained annual (1970–2023) numbers of adult salmon catches (coastal fishery and hatchery broodstock) and hatchery chum salmon fry releases from the Hokkaido Salmon Enhancement Association (HSEA, 2024), Fisheries Research and Education Agency (FRA, 2024), and the Aomori Prefectural Government (APG, 2024). FRA (2023) biological and reproductive trait data included time series of fry size-at-release and release month. We selected hatchery release events with fry body weight (BW) and fork length (FL) data (1998–2020, Hokkaido: 184 rivers, 37,886 release events, *n* = 21,332,093 fish; Honshu: 141 rivers, 19,605 release events, *n* = 9,588,413 fish). The data also included adult age composition at return (1997–2020, Hokkaido: 52 rivers; Honshu: 102 rivers; 1,902 surveys, *n* = 1,210,717 fish). Adult salmon trait data included egg weight, fecundity, FL, age-at-return, and gonad weight (1997–2020, Hokkaido: 10 rivers; Honshu: 14 rivers; *n* = 37,533 fish), and adult age-at-return (counts of annular marks on fish scales), including six age groups: age-2 (*n* = 25), age-3 (*n* = 2,892), age-4 (*n* = 20,643), age-5 (*n* = 13,118), age-6 (*n* = 846), and age-7 (*n* = 9). We used trait data such as FL, egg weight, and fecundity of age-4 and age-5 fish for our analysis.

We obtained annual commercial catch data for all marine species in Japan (1980–2020) from the Japan Ministry of Agriculture, Forestry and Fisheries (MAFF, 2023). Catch statistics (1998– 2022) for Canada, Russia, and USA pink and chum salmon were available from the North Pacific Anadromous Fish Commission (NPAFC, 2023). For SST analyses, we downloaded Japan coastal 0.25°×0.25° daily mean SST data (1982–2020) from Hokkaido (18 areas) and Honshu (12 areas) from the Japan Meteorological Agency (JMA, 2022) and global 0.25°×0.25° monthly mean SST data (1982 to 2022) from the NOAA Optimum Interpolation (OI) SST V2 high-resolution dataset (NOAA, 2023a).

### 2.3 Catch, release, and reproductive traits

We summarized Japanese chum salmon catch, release, and reproductive traits data by seven management regions in northern Honshu and Hokkaido, Japan (Figure 1a). To estimate the distribution of release times in each region, we counted the number of release events by month. We also calculated the mean values of size-at-release, biological traits, and age composition by management region. Since the number of individuals released (𝑛_𝑘_) varied among release events (*k*=1,…,*K*), we calculated the weighted mean size-at-release for BW (g) (1998–2020) as 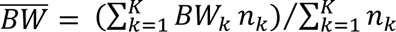. Here, the suffix for a region was omitted for simplicity. We calculated the mean values of the traits and the relative fecundity (eggs/*FL*(cm)^3^ × 10^3^) of the age-4 and age-5 fish from 1997 to 2020. The weighted mean proportion of age-*a* fish was calculated as 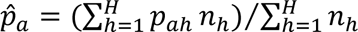, where *h* refers to each survey, 𝑝_𝑎ℎ_ is the proportion of age-*a* fish in survey *h*, 𝑛_ℎ_ is the sample size (individuals), and *a* refers to age at return (*a* = 2,…,8).

### 2.4 Adult return rates (marine survival index)

We assumed all adult chum salmon returning to Japan were hatchery reared (see Discussion, section 4.1), and adult return rates are survival rates from release to return. In this study, we defined the adult return rate as an index of the marine survival of chum salmon (James et al. 2023). Hatchery programs have been managed in five management regions in Hokkaido and by prefecture in Honshu, and previous studies evaluated return rates by management region (Saito & Nagasawa, 2009; Miyakoshi et al., 2013; Kitada, 2014). The management regions coincide with the genetic structure of the population (Beacham et al., 2008). We estimated return rates for the current hatchery management regions (Figure 1a). We estimated the total number of fish returning from releases in year *t* (1998–2019) (𝑌_*t*_, coastal commercial catch plus river catch for hatchery broodstock) by

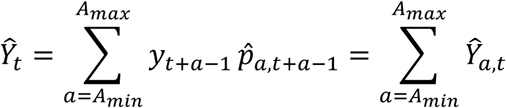

where 𝑦_*t*+𝑎−1_ is the observed number of fish returned in year *t* + 𝑎 − 1 from fry released in year *t*, and 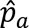 is the estimated proportion of age-*a* fish among the fish returned in year *t* + 𝑎 − 1. We estimated the return rate of fry released in year *t* by 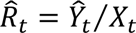 is the number of hatchery-reared fry released in year *t*. We estimated the age-specific return rate as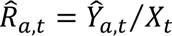. We summed the number of fish returned from ages 2–5 (𝐴*_min_* = 2, 𝐴*_max_* = 5). Most returns were ages 2–5 fish, and few returns were age-6. There were no age-7 and age-8 fish in returns to any region except Hokkaido Okhotsk, where returns of these age groups were very low. We estimated the age composition of returning adult chum salmon from available (1997–2020) data (Figure S1). We estimated the age composition in 2021–2023, where no data were available, as a moving average of the previous three years to account for the recent trend of increasing numbers of young fish. Based on age composition, we calculated the number of returning fish (ages 2–5) for each release year. Then, we estimated return rates and age-specific return rates from releases in 1998–2019. We measured distances between return rates in the management regions by 1 − *r_uv_* and illustrated distances by a neighbor-joining (NJ) tree (Saitou & Nei, 1987) using the “MEGA11” package (Tamura et al., 2021), where *r_uv_* is the correlation coefficient between pairs of return rate time series in regions *u* and *v*.

To assess and visualize continuous changes in log-transformed return rates and covariates over the years, we fit GAMs to the data using the ‘gam’ function in the “mgcv” package (Wood, 2017) in R (R Core Team, 2024), assuming normal errors. Year was included as a nonlinear smoothed term (explanatory variable) because these covariates varied across years. The trend of the traits over the years is captured by the non-linear and/or linear year effects (Wood, 2004, 2011).

### 2.5 Long-term changes in chum salmon thermal habitat

To understand long-term changes in SST in the chum salmon habitats covered by our regression model (see section 2.8), we mapped monthly mean 1°×1° SST isoclines in the North Pacific Ocean for 1990, 2000, 2010, and 2022, using the Plot COBE Sea Surface Temperature system (NOAA, 2023b), based on the COBE2 SST dataset from JMA. The 1°×1° monthly mean SST isoclines around Japan for 2000, 2010, and 2022 were generated similarly. We mapped the available temperature range of chum salmon (4–13°C), based on previous predictions: optimal growth and feeding (8–12°C), available habitat for feeding (5–13°C), and overwintering temperature range (4–6°C) (Kaeriyama et al., 2012; Urawa et al., 2018). Langan et al. (2024) found that chum salmon in the North Pacific Ocean were frequently caught at SSTs below 4°C in spring and summer. Their analysis was based on unknown population mixtures of chum salmon in historical high-seas research vessel catches. It is quite possible that southern populations such as Japanese chum salmon are less cold tolerant than northern populations.

### 2.6 Japanese chum salmon migration route and SST

We calculated monthly mean SSTs for 30 coastal areas in Japan (JMA, 2022). From these, we calculated monthly mean SSTs for the seven management regions (Figure 1a), and for release periods of chum salmon fry (Figure S2): Hokkaido Okhotsk (Apr–Jun), Hokkaido Nemuro (Mar–Jun), Hokkaido Pacific East (Mar–Jun), Hokkaido Pacific West (Mar–May), Hokkaido Sea of Japan (Mar–May), Honshu Pacific (Feb–May), and Honshu Sea of Japan (Feb–Apr).

Chum salmon fry born in rivers along the Pacific coast of Japan migrate to the Sea of Okhotsk (Honda et al., 2017, 2025). Juvenile chum salmon from all regions of Japan are found in the Sea of Okhotsk, and the abundance of juveniles is significantly correlated (positive) to adult returns to Japan (Urawa & Bugaev, 2021). Juvenile chum salmon were abundant in coastal areas of the Sea of Okhotsk from May to June when SST was between 8 and 13°C, and disappeared from coastal waters after late June when SST exceeded 13°C (Nagata et al., 2007). Based on these results, we assumed that chum fry released in all areas would migrate to the Sea of Okhotsk and disperse offshore in July (Figure 1a).

We identified the locations of four main offshore and high-seas areas of Japanese chum salmon distribution (Figure 1b) based on information from previous studies (Myers et al., 2007; 2016; Nagata et al., 2007; Urawa et al., 2018, 2022): 1. Southern Sea of Okhotsk (45.2°N–46.5°N, 142.5°E–148.5°E), 2. Western North Pacific (42°N–45°N, 158°E–165°E), 3. Central Bering Sea (53°N–58°N, 175°E–170°W), 4. Central Gulf of Alaska (48°N–56°N, 140°W–150°W). Using “sst.mnmean.nc” data (NOAA, 2023a), we calculated the monthly mean SSTs (1982– 2022) for salmon distribution areas (Figure 1b) using the ‘fldmean’ function in the “cdo-2.1.1” package (Max Planck Institute, 2023). We mapped areas and migration routes using the “sf” package in R. We used monthly mean SSTs (1998–2021) in our regression model.

### 2.7 Potential predators of chum salmon fry

We expected that climate-driven changes in abundance and distribution of piscivorous marine fish predators are related to decreases in the marine survival of Japanese chum salmon fry in coastal waters. We used principal components analysis (PCA) to reduce a large number of potential predator variables to a smaller number that explained most of the variance in the predator data. To characterize temporal changes in abundance of marine fish for the first stage of screening for potential predators, we performed PCA on the annual commercial catch data for the marine species in 1980-2020 (MAFF, 2023) using the ‘prcomp’ function in R. For the second stage of screening for potential predators, we visualized trends in catches of the marine species analyzed in PCA. Because catches vary by geographic area and season, we obtained catch data of potential marine fish predator species identified by PCA by reporting area in Hokkaido and by prefecture in Honshu (MAFF, 2023), which should reflect their fishing seasons and migration patterns. Commercial catches of potential marine predators identified by PCA and chum salmon were mapped in the coastal range of Japanese chum salmon to illustrate decadal-scale shifts in predator-prey abundance and distribution.

### 2.8 Regression analysis of adult return rates

Our regression model describes the relationship between return rates of chum salmon fry released in year *t* across all ages 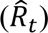 and explanatory variables (*i*=1,…,*s*) such as SSTs, size-at-release, and catch of potential predators and competitors. We transformed the return rate to the log scale for model fitting in a multiple linear regression:

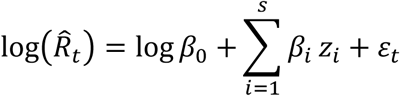

where 𝛽_0_ is the intercept, and the explanatory variables are denoted by 𝒛. The regression coefficients were denoted by 𝜷 and 𝜀_*t*_∼𝑁(0, 𝜎^2^) is the error that we assumed to be independently and normally distributed. We did not include interactions between explanatory variables. We selected the explanatory variables by AICc (Sugiura, 1978), which is the small sample correction of the Akaike Information Criterion (AIC, Akaike, 1973).

We estimated regression coefficients based on standardized explanatory variables, which allow for consistent comparison of different explanatory variables across management regions as done in salmon studies (e.g., Oke et al., 2020; Ohlberger et al., 2022). The use of an unstandardized measure (regression coefficient) is preferred to a standardized measure if the units of measurement (e.g., fry size-at-release, SST, and abundance of competitors) are used to describe effect sizes at a “practical” or “real world” level (Wilkinson et al., 1999). Therefore, we also estimated the regression coefficients using unstandardized explanatory variables, as done in Malick et al. (2023). In the latter, we define effect size as the effect of the unit change of an explanatory variable on the change in return rate. The effect size and 95% confidence intervals were calculated by 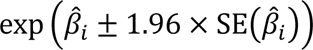 for coefficient *i*. We predicted return rates by region using the ‘predict’ function in R based on regression coefficients using the unstandardized measure and compared them to the observed return rates.

When many explanatory variables are included, the model selection procedure may select an inappropriate model as the best model due to complex multicollinearity. To avoid this problem, we created three subset models of stressors (see below), performed variable selection within each subset model, and compared the best models for combinations of the subset models and the full model, which included all subset models. In each model, we used the variance inflation factor (VIF) to diagnose multicollinearity among explanatory variables. We determined variables that could be included in a model by sequentially removing each explanatory variable with the highest VIF value until all remaining VIFs were less than the threshold of 3.0 (Zuur et al., 2010), using the ‘VIF’ function in the “car” package in R. Our model selection procedure examined all combinations of explanatory variables (2*^s^*, in our case *s* = 17 without subset models). Analysis using subset models and a strict VIF threshold should help reduce the risk of selecting incorrect models due to multicollinearity.

The dependent variable is the adult return rates of fry released in 1998–2019. We used 17 hypothesized stressors (covariates) for explanatory variables (*s* = 17) and grouped them into three broad categories of subset models: (1) hatchery carryover (three metrics), (2) ocean condition (five metrics), and (3) predators and competitors (nine metrics) (Table 1). The hypothesized hatchery carryover covariates included relative fecundity and egg size of females in year *t*-1, where *t* is the year of fry release, and fry size-at-release. Ocean-condition covariates included SSTs at fry release and in Japanese coastal management regions, summer SSTs in the southern Sea of Okhotsk and central Bering Sea, and winter SSTs in the western North Pacific and central Gulf of Alaska (see Figure 1). Predator and competitor covariates included predation in coastal Japan regions and competition in high-seas areas (Table 1). We identified yellowtail (*Seriola quinqueradiata*), also known as Japanese amberjack, as a potential predator of chum salmon fry (see section 3.4). Age-4 chum salmon were the dominant age group in adult returns to Japan (Figure S1). We used the annual commercial salmon catch (total number of fish) of chum salmon in Russia, the United States, and Canada in year *t*+3 as indices of intraspecific competition with Japanese chum salmon released in year *t*, as they return in year *t*+3 as age-4 fish. The correlation between chum and pink salmon catches in Russia and US/Canada was very weak (*r* = 0.0005 for chum and *r* = −0.18 for pink), and we used Russia and US/Canada catches as covariates. Chum salmon were assumed to spend three years in the high seas and to compete when sharing the same environment. Pink salmon have a fixed life span of two years (Quinn, 2018). We used annual commercial catches in Russia and North America (US and Canada, combined totals) in years *t*+1, *t*+2, and *t*+3 as indices of interspecific competition with Japanese chum salmon released in year *t*. The ocean distribution of chum and pink salmon overlaps throughout the North Pacific (Langan et al., 2024). We assumed cohorts of pink salmon compete with chum during the first (*t*+1), second (*t*+2), and third (*t*+3) ocean years.

We performed regression analysis independently for each subset model and combinations of subset models (including the full model) with explanatory variables less than 3.0 VIF (see, Table S2). This procedure reduces the number of parameters to be estimated and increases the information available for inference, resulting in robust inference. In each analysis, we selected the most parsimonious model (hereafter, the best model) with the minimum AICc using the “MuMIn” package in R. We compared all models by their minimum AICc values. To summarize the results, we presented the regression coefficients of the best model for the selected subset models. We expected there may be differences in covariate effects among management regions, probably due to differences in the quality and size of fry at release and the coastal environments. Therefore, we modeled the management regions separately (see section 3.1). In three of the four regions, the minimum of the minimum AICc values was achieved by multiple hierarchical subset models. In such cases, we chose the most parsimonious subset models. The estimated regression coefficients were compared with the results obtained by model averaging over all models with ΔAICc values less than 2 (ΔAIC < 2), which suggests substantial support for the model (Burnham & Anderson, 2002). Practical model averaging provides the weighted averages of the regression coefficients over the models. The weights represent the contributions of the models and are proportional to exp (− 1⁄2 × AICc). All data and the R script used for the regression analysis are available (see data availability statement).

## 3| RESULTS

### 3.1 Adult return rate (marine survival index)

For Japan as a whole, the number of hatchery-released fry increased until the early 1980s, reaching 2 billion, and then decreased to about 1.3 billion at present (Figure 2). In the Honshu Pacific, there has been a steady decline in releases since the mid-1990s and a rapid decline since 2021. Decreasing numbers of released fry were also observed in Hokkaido Pacific East and Honshu Sea of Japan, while stable numbers were observed in Hokkaido Nemuro, Hokkaido Pacific West, and Hokkaido Sea of Japan. Only in Hokkaido Okhotsk did the number of fry released increase slightly but continuously. Releases were largest in Honshu Pacific, while other regions had similar release levels.

**FIGURE 2.**
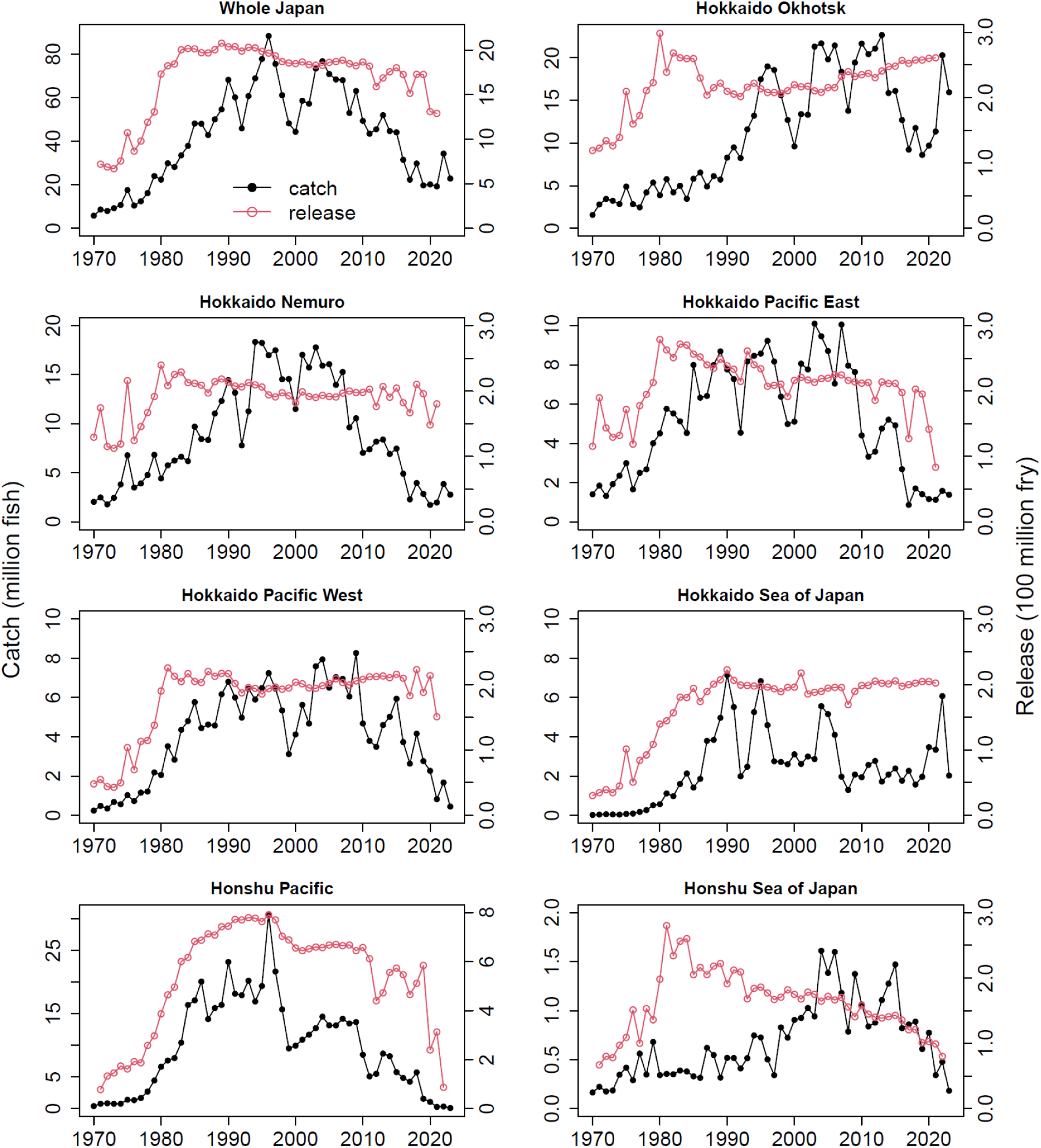
Number of hatchery chum salmon fry released (red line) and adult returns (coastal catch plus hatchery broodstock catch; black line) in seven management regions in Japan (1970– 2023; Figure 1a).

The total number of adult chum salmon returns throughout Japan increased rapidly until 1996 (88 million fish), and then declined to the current level of 20 million fish (Figure 2). The highest regional returns were in the Honshu Pacific in 1996 (31 million), and then declined continuously to the historical regional minimum (∼10 thousand). Among other regions, adult returns were highest in northern Hokkaido regions (Okhotsk and Nemuro) and lowest in Sea of Japan regions (Hokkaido and Honshu). Since approximately 2008, adult returns decreased in all management regions except the Hokkaido Okhotsk and Hokkaido Sea of Japan, and the decrease was most severe in the Hokkaido and Honshu Pacific regions.

The mean adult return rate with the interquartile range of the 5th and 95th percentiles (including ages 2–5) for 1998–2019 releases was highest in Hokkaido Okhotsk (6.9 [3.0, 10.4]%), followed by Hokkaido Nemuro (4.5 [1.1, 9.5]%), Hokkaido Pacific East (2.3 [0.6, 5.5]%) and Hokkaido Pacific West (2.3 [0.6, 4.5]%), Hokkaido Sea of Japan (1.4 [0.5, 2.9]%), Honshu Pacific (1.2 [0.1, 2.3]%), and Honshu Sea of Japan (0.6 [0.4, 1.4]%) (Figure S3). The GAMs showed that log-transformed return rates decreased over time in all management regions, except in the Hokkaido Okhotsk and the Hokkaido Sea of Japan, where return rates increased recently (Figure S4). The nonlinear and/or linear year effect was significant in all management regions except in the Hokkaido Sea of Japan and the Honshu Sea of Japan (Table S1).

The NJ tree analysis showed that the Hokkaido Okhotsk, Honshu Sea of Japan, and Hokkaido Sea of Japan adult return rates were distinct, and the Hokkaido Nemuro, Hokkaido Pacific East and West, and Honshu Pacific return rates were similar (Figure 3). As a result, we used four coastal regions in Japan in our regression analysis: (1) Hokkaido Okhotsk, (2) Hokkaido and Honshu Pacific (combined returns to Hokkaido Nemuro, Hokkaido Pacific East and West, and Honshu Pacific), (3) Hokkaido Sea of Japan, and (4) Honshu Sea of Japan.

**FIGURE 3.**
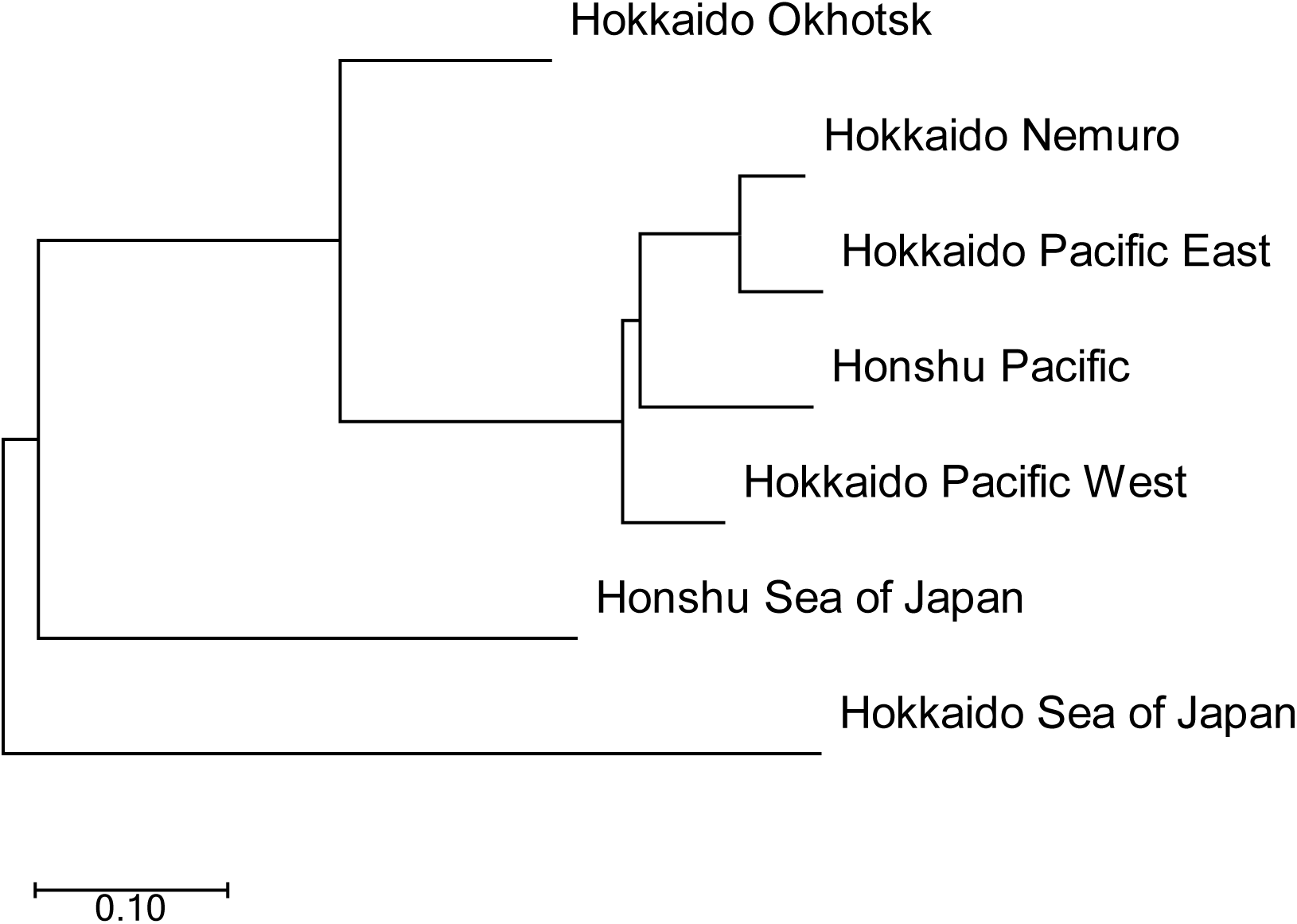
Neighbor-joining tree based on correlation coefficients of chum salmon adult return rates between pairs of seven management regions in Japan, 1998–2019 (Figure 1a).

### 3.2 Long-term changes in chum salmon thermal habitat

Decadal-scale maps of SST isoclines showed that until 2000, the summer and winter thermal habitat of chum salmon was stable (Figure 4). After 2010, the optimal summer habitat for chum salmon growth and feeding (especially, 8–11 °C isotherms) shifted north and contracted longitudinally, and the winter habitat remained relatively stable. Evaluation of intra-seasonal changes in spring and summer thermal habitat revealed a progressive northward shift in optimal habitat, from a slight shift in early spring (March and April) to a larger shift in early summer (May and June), northward longitudinal contraction in late summer (August and September), and a slight northward shift in fall (October and November) (Figures S5–S8). In coastal Japan, the optimal thermal habitat for chum salmon fry migrating to the Sea of Okhotsk shifts northward in spring and early summer (April–June), especially in the Sea of Japan (Figure S9).

**FIGURE 4.**
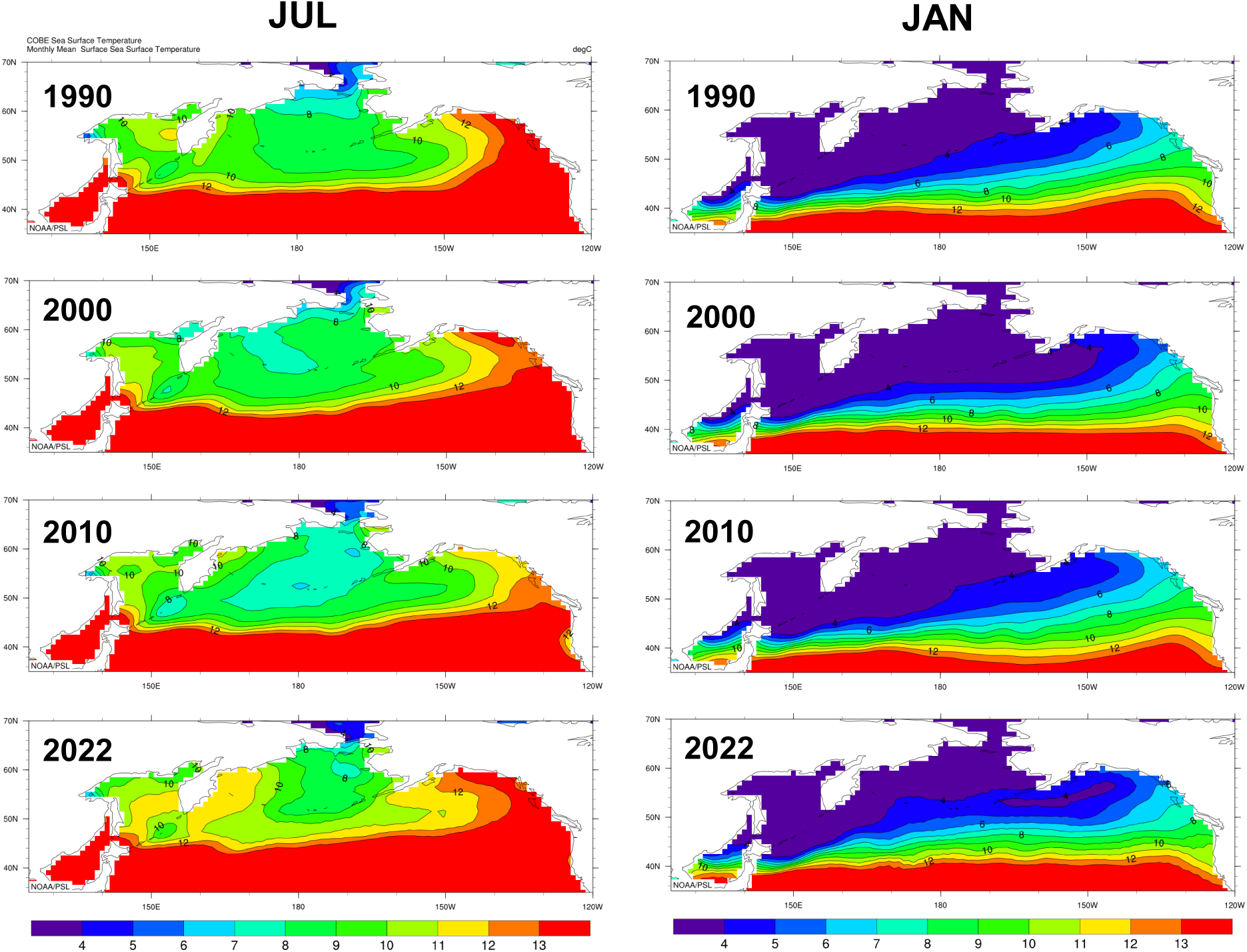
Sea surface temperature (SST) isotherms in recent decades (1990-2022) showed a northward shift and longitudinal contraction of chum salmon summer (July) thermal habitat in the North Pacific Ocean, while winter (January) habitat was relatively stable. Blue, green and orange show the range of Japanese chum salmon thermal limits (4–13°C): optimal growth and feeding habitat (8–12°C), swimming and feeding habitat (5–13°C), and overwintering habitat (4–6°C) (Kaeriyama et al., 2012; Urawa et al., 2018). Red (>13°C) and purple (<4°C) indicate critical habitat, lost to ocean production of Japanese chum salmon.

### 3.3 Japanese chum salmon migration route and SST

In the Hokkaido Nemuro region, the mean SST increased significantly, while the Hokkaido Pacific West and Honshu Pacific regions showed a similar pattern; the mean SST decreased until around 2010 and then slightly increased (Figure S10, Table S1). In the other regions, the year effects were insignificant.

The summer and winter SSTs used in our regression analysis (1998–2022) as common explanatory variables for all offshore (southern Sea of Okhotsk) and high-seas areas (Figure 1b) were southern Sea of Okhotsk July SST, western North Pacific January–April SST, central Bering Sea August–September SST, and central Gulf of Alaska January–April SST (Figure S11a). A GAM (Figure S11b) showed that July SST in the southern Sea of Okhotsk (range, 10.8–14.2°C) increased after the early 2000s, and a strong increase was observed after around 2015. In the western North Pacific, winter SST in January–April (range, 4.6–6.1°C) increased after around 2015. In the central Bering Sea, August–September SST (range, 8.3–11.4°C) continued to increase since the late 1990s but decreased after around 2015. In the central Gulf of Alaska, a strong increasing trend in January–April SST (range, 4.6–7.2°C) was observed after around 2010, but SST decreased after around 2015. Similar trends were observed in the western and eastern North Pacific.

### 3.4 Potential predators of chum salmon fry

We summarized the landed commercial catches of all marine species in Japan with no missing data from 1980 to 2020 (56 species). We excluded marine mammals because catches are subject to strict limits, and catches were not recorded by species but by total marine mammals. We also excluded species known not to be predators of salmon (e.g., sardine, smelt, algae, shellfish, and crustaceans). This resulted in catch data for 30 fish species. The PCA biplot identified differences in commercial fishery catch levels as the primary component (PC1, 46% variance), and the second component (PC2, 18% variance) corresponded to catch trends before and after 1998 (Figure 5a). The PCA confirmed that changes in commercial catches of 30 marine fishes in Japan after 1998 were largely characterized by two species that had positive PC1 values and increased in recent decades: yellowtail and Japanese Spanish mackerel (*Scomberomorus niphonius*). We visualized trends in catches of the 30 species (1980–2020) and confirmed that catches increased after 1998 only in the two species identified by the PCA analysis (unpublished figure, S. Kitada). Yellowtail and Japanese Spanish mackerel are strong piscivores. There was a northward shift of yellowtail landings into the range of Japanese chum salmon (Figure 5b). In contrast, increased landings of Japanese Spanish mackerel occurred only in the eastern area of Japan, with slight increases in the Honshu North Pacific area. Based on these results, we identified yellowtail as a possible new predator of chum salmon fry.

**FIGURE 5.**
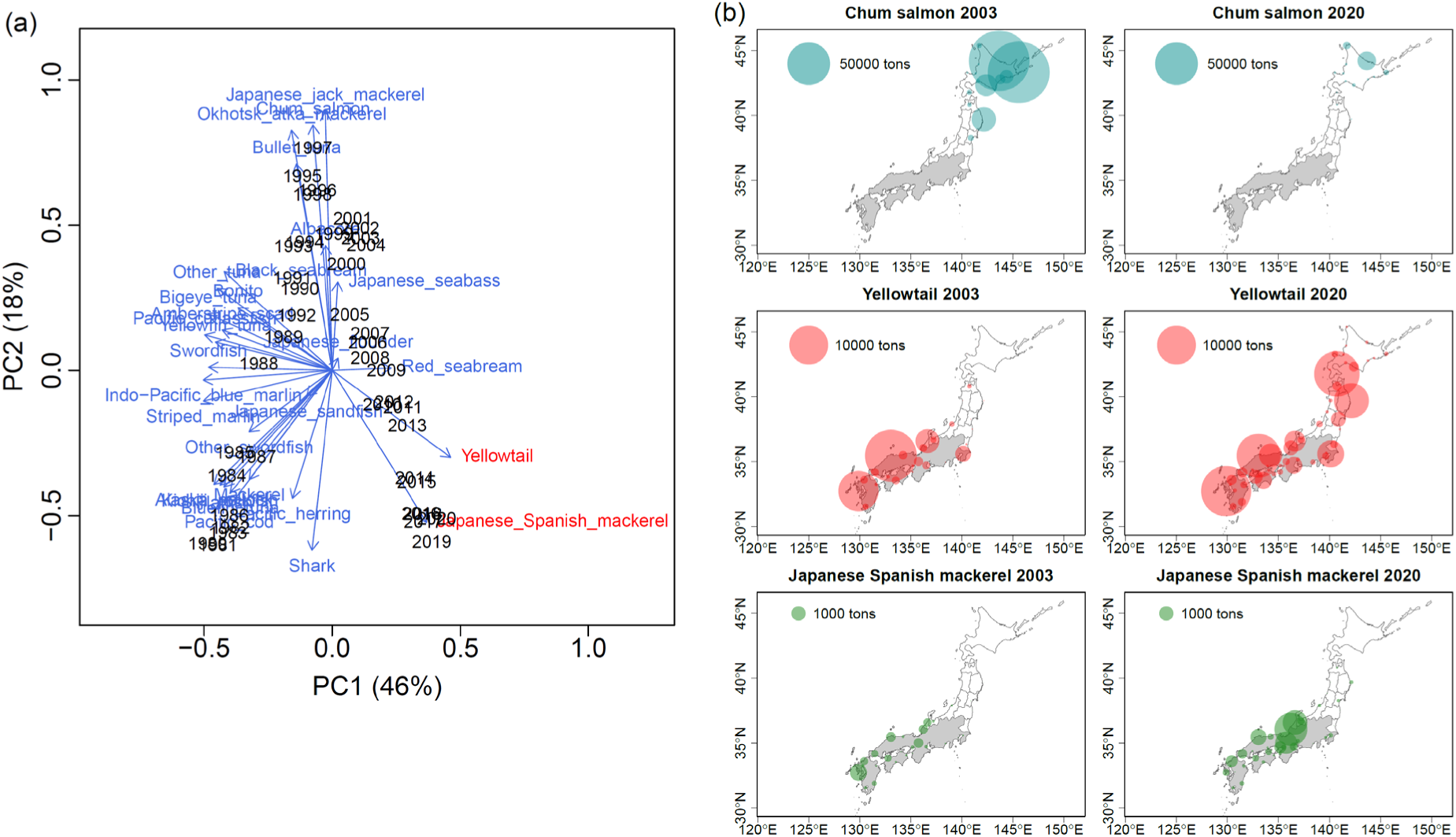
Northward shifts of two marine piscivorous fishes in Japan. (a) Principal component analysis biplot based on catch statistics of 30 marine fish species (1980 - 2020) showed increasing catches of yellowtail and Japanese Spanish mackerel. (b) Two-decadal shifts in commercial catches of chum salmon and two piscivores.

### 3.5 Regression analysis of adult return rates

All models used explanatory variables with a VIF less than the threshold of 3.0. Model selection results for all subset models are shown in Table S2. Egg size and relative fecundity of age-4 and age-5 fish were examined, but these explanatory variables were selected only for Hokkaido and Honshu Pacific, where age-4 fish had a smaller AICc (30.0) and larger *R*^2^ (0.79) than age-5 fish (AICc = 38.2 and *R*^2^ = 0.61). Therefore, we used egg size and relative fecundity of age-4 fish in our regression analysis. The full model results and the best subset model results were the same for all management regions except for the Hokkaido Sea of Japan (Table 2; Tables S2 and S3). The results of the model averaging (Table S4) were qualitatively the same as the best models (Table 2) for all regions. The explanatory variables selected in the best models were region-specific or common to some areas, or both.

**TABLE 2.**
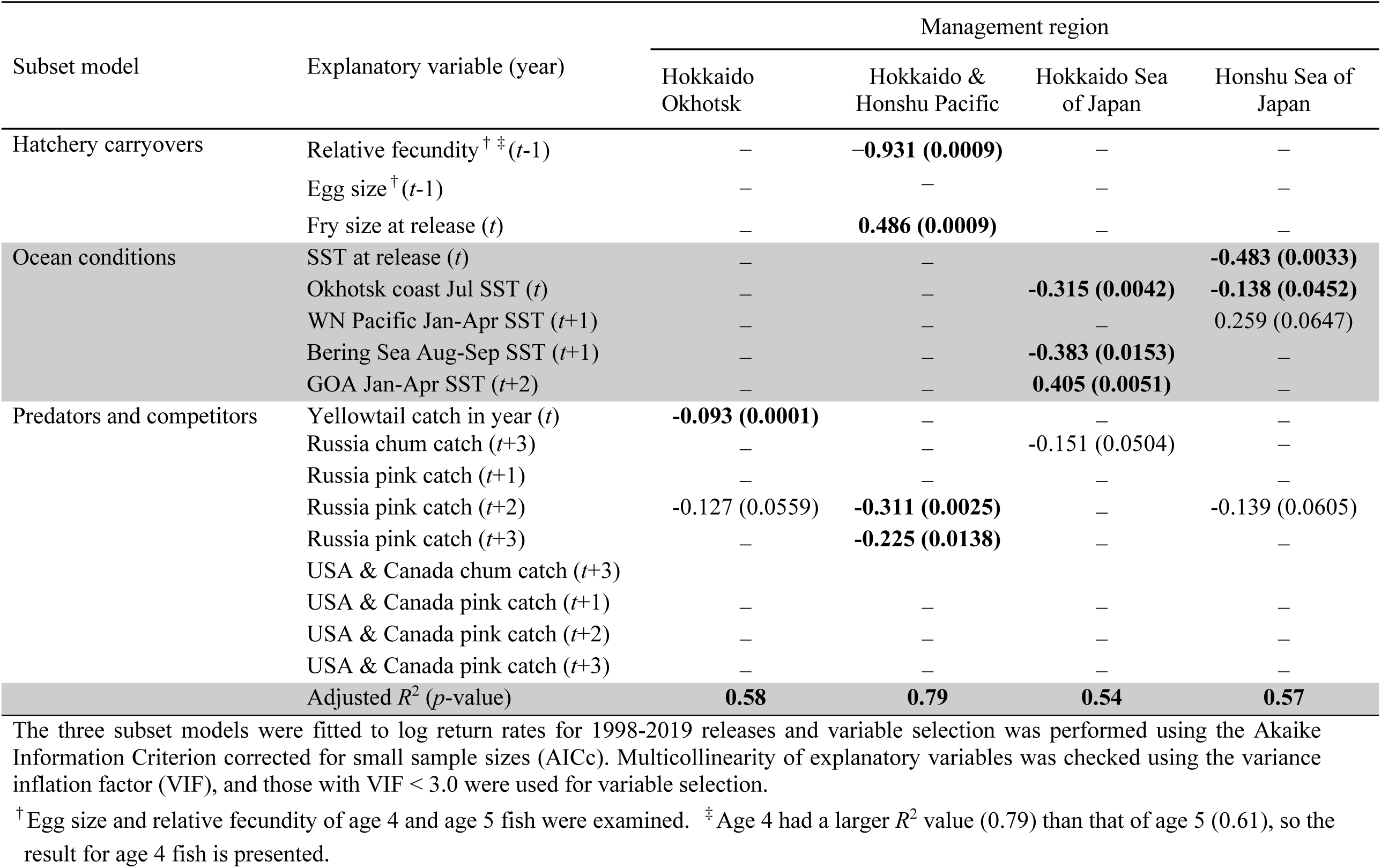
Best regression models of Japanese chum salmon survival in four coastal regions in Japan and regression coefficients with their *p*-values in parentheses. Bold indicates significant relationship (𝛼 = 0.05). *t* = hatchery release year of chum salmon fry. Sea surface temperature (SST); Western North Pacific (WN Pacific); Gulf of Alaska (GOA). See Figure 1a for management region locations.

For ocean conditions, the effects of SST were found only for fry released in the Hokkaido Sea of Japan and Honshu Sea of Japan regions (Table 2). SST at release (year *t*) and summer SST (July) in the Sea of Okhotsk (year *t*) and the Bering Sea (August-September) (year *t*+1) were negatively associated with Japanese chum salmon return rates. In contrast, winter SST (January-April) in the western North Pacific (year *t*+1) and the Gulf of Alaska (year *t*+2) were positively associated with return rates. For predators and competitors, we found negative associations between return rates and yellowtail catch in the year of fry release only in the Hokkaido Okhotsk region, while negative effects of Russian pink and/or chum salmon catches were found in all regions. Russian chum salmon catch in year *t*+3, and Russian pink salmon (adult, age-1) catch in years *t*+2 and *t*+3 were negative. No associations between Japanese chum salmon return rates and US and Canadian chum and pink salmon catches were found. For hatchery carryovers, fry size-at-release was positively associated with salmon return rates and relative fecundity was negatively associated with return rate only in the Hokkaido and Honshu Pacific region.

The GAMs indicated a similar trend in fry size-at-release in Hokkaido Okhotsk, Hokkaido Sea of Japan, and Honshu Pacific, where fry size-at-release increased or remained at similar levels until around 2010 and then declined (Figure S12, Table S1). The decline in fry size-at-release was remarkable in the Honshu Pacific and Honshu Sea of Japan regions. In contrast, the fry size-at-release increased after about 2015 in Hokkaido Nemuro. Fry size-at-release varied among regions (Figure S13) and the mean was smallest in the Hokkaido Sea of Japan, while the Honshu Sea of Japan had the greatest variation with a recent minimum.

We found that the relative fecundity of parent females was negatively associated with return rates in the Honshu Pacific region. Relative fecundity was negatively correlated with egg size in all management regions except in the Hokkaido Sea of Japan region (unpublished figure, S. Kitada). GAMs confirmed that egg size for age-4 fish has decreased over the last two decades in all regions, except Hokkaido Okhotsk (Figure 6, Table S1). The trend was significant in all regions, except Honshu Sea of Japan. Similar decreasing trends in egg size for age-5 fish were also observed (Figure S14, Table S1). Egg size varied among the management regions, and was smallest in the Hokkaido Sea of Japan region (Figure S15).

**FIGURE 6.**
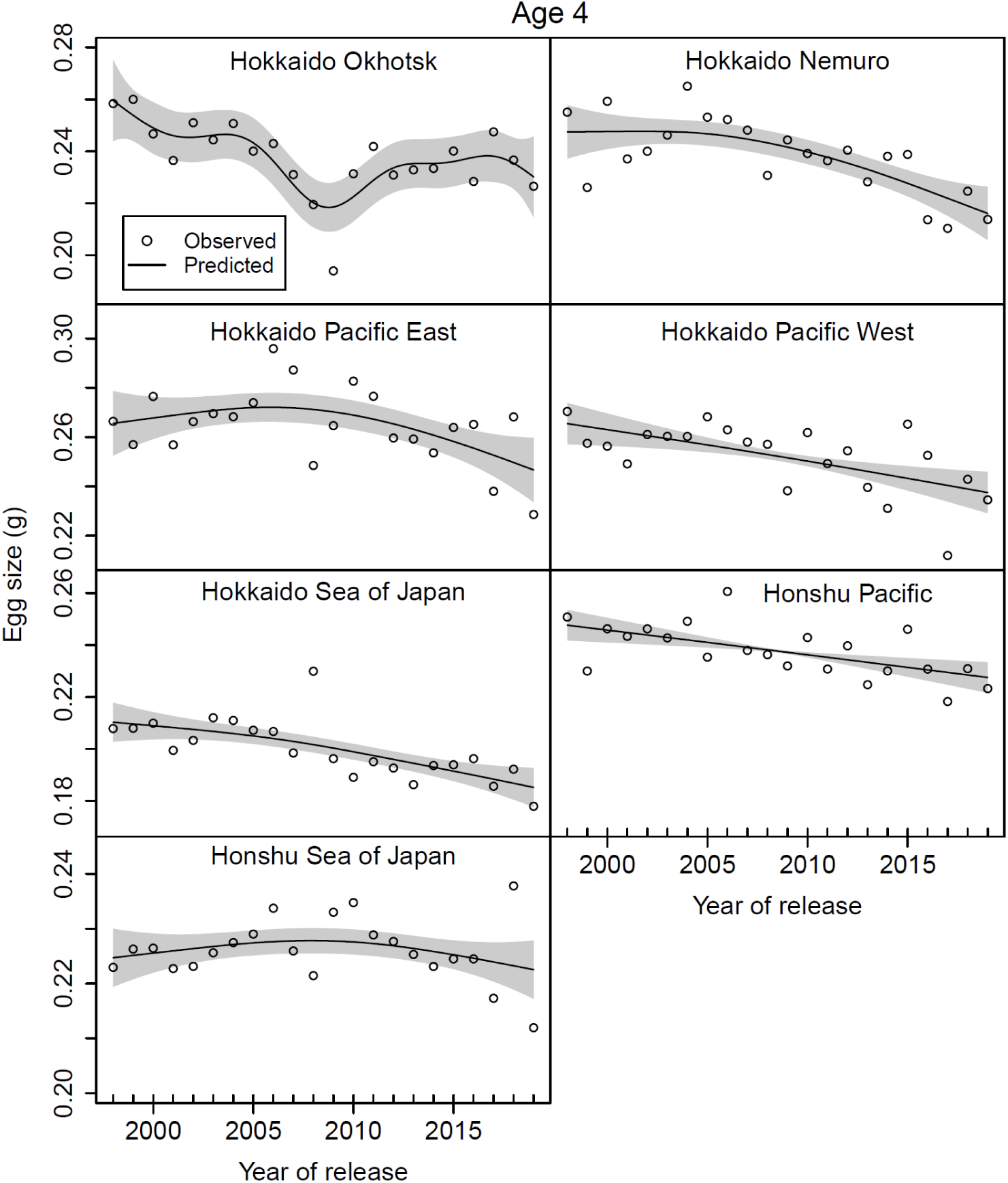
Egg size of age–4 chum salmon has generally decreased in all regions, except Hokkaido Okhotsk as shown by the nonlinear and/or linear year effect from GAMs (1998-2019). The trend was significant in all regions, except Honshu Sea of Japan (Table S1). Open circles show observed values. The black lines show predicted values with shaded areas of 2×standard errors.

The standardized model effect size of explanatory variables on adult returns allowed comparison of the magnitude of the effects across the four regions (Figure 7). The largest positive effect was fry size-at-release (year *t*), which was equal to winter SST (January–April) in the Gulf of Alaska (year *t*+2), followed by winter SST (January–April) in the western North Pacific (year *t*+1). On the other hand, parental relative fecundity (year *t*-1) had the largest negative effect, followed by Russia pink and/or chum salmon catches which varied among regions. The negative effect of summer SST in the Bering Sea (August–September) (year *t*+1) was the second largest, followed by summer SST in the Sea of Okhotsk (July) (year *t*) and SST at the time of release (year *t*). The negative effect of yellowtail (year *t*) was similar to summer SST in the Bering Sea.

**FIGURE 7.**
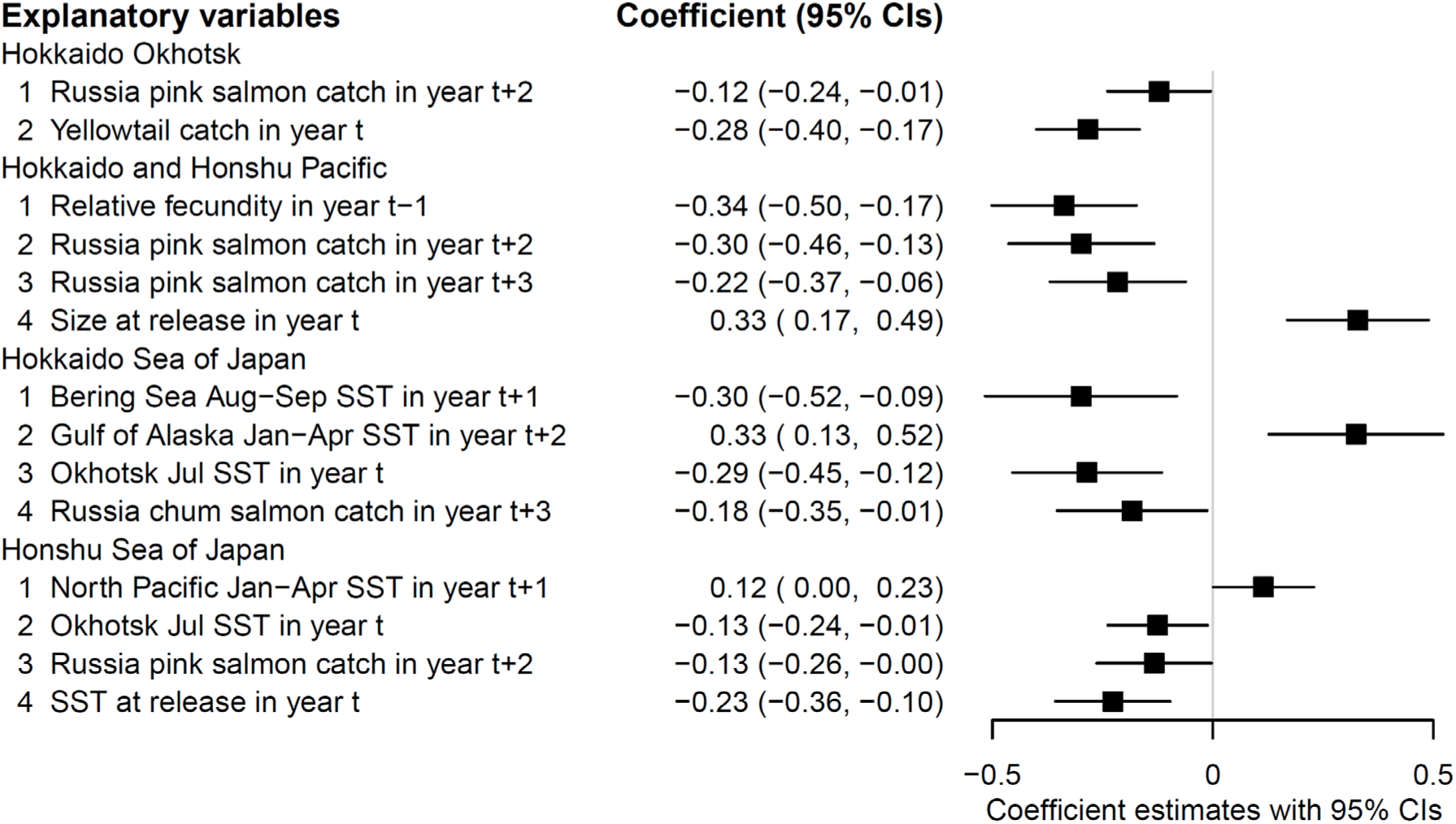
Standardized effects of hypothesized explanatory variables on Japanese chum salmon marine survival (adult return rates across regions). Regression coefficients of best regression models with 95% confidence intervals in four coastal regions in Japan, estimated using standardized explanatory variables. *t* = hatchery release year of chum salmon fry. Sea surface temperature (SST). See Figure 1a for management region locations.

The practical effect size in a regression model describes how much of a change in the dependent variable is expected for a given change in the independent variable in each region. The effect size for each regression coefficient, measured by 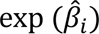, represents the rate of change per practical unit in each management region (Figure S16), which showed a similar pattern to the standardized effects of the explanatory variables (Figure 7). The area-specific practical effect sizes (exp(0.486) = 1.625) showed that a 0.1 g increase in hatchery fry size increased the return rate by 62.5% ((1.625-1)×100) in the Hokkaido and Honshu Pacific region (Table 3).

**TABLE 3.** Regression coefficients and practical effect sizes on Japanese chum salmon return rates in four coastal regions in Japan. *t* = hatchery release year of chum salmon fry. Sea surface temperature (SST); Western North Pacific (WN Pacific); Gulf of Alaska (GOA).

| Region | Explanatory variable (year) | Estimate<br>$\hat{\beta}_i$ | Standard<br>error | Effect<br>size <sup>†</sup> | 95%<br>lower | 95%<br>upper | Unit | Practical<br>effect (%) <sup>‡</sup> |
| --- | --- | --- | --- | --- | --- | --- | --- | --- |
| Sea of Okhotsk | Russia pink catch ($t+2$) | -0.127 | 0.062 | 0.881 | 0.779 | 0.995 | 100 million fish | -11.9 |
| | Yellowtail catch ($t$) | -0.093 | 0.020 | 0.911 | 0.877 | 0.947 | 100 tons | -8.9 |
| Hokkaido and<br>Honshu Pacific | Relative fecundity ($t-1$) | -0.931 | 0.232 | 0.394 | 0.250 | 0.620 | 100 eggs <sup>§</sup> | -60.6 |
| | Russia pink catch ($t+2$) | -0.311 | 0.088 | 0.732 | 0.617 | 0.870 | 100 million fish | -26.8 |
| | Russia pink catch ($t+3$) | -0.225 | 0.082 | 0.799 | 0.680 | 0.938 | 100 million fish | -20.1 |
| | Fry size at release ($t$) | 0.486 | 0.121 | 1.625 | 1.283 | 2.059 | 0.1g | 62.5 |
| Hokkaido Sea of<br>Japan | Bering Sea Aug-Sep SST ($t+1$) | -0.383 | 0.142 | 0.682 | 0.516 | 0.901 | 1°C | -31.8 |
| | GOA Jan-Apr SST ($t+2$) | 0.405 | 0.126 | 1.499 | 1.171 | 1.918 | 1°C | 49.9 |
| | Okhotsk coast Jul SST ($t$) | -0.315 | 0.096 | 0.730 | 0.605 | 0.880 | 1°C | -27.0 |
| | Russia chum catch ($t+3$) | -0.151 | 0.072 | 0.859 | 0.746 | 0.990 | 10 million fish | -14.1 |
| Honshu Sea of<br>Japan | WN Pacific Jan-Apr SST ($t+1$) | 0.259 | 0.131 | 1.295 | 1.002 | 1.675 | 1°C | 29.5 |
| | Okhotsk coast Jul SST ($t$) | -0.138 | 0.064 | 0.871 | 0.769 | 0.987 | 1°C | -12.9 |
| | Russia pink catch ($t+2$) | -0.139 | 0.069 | 0.870 | 0.760 | 0.997 | 100 million fish | -13.0 |
| | SST at release ($t$) | -0.483 | 0.142 | 0.617 | 0.467 | 0.814 | 1°C | -38.3 |

<sup>†</sup> $\exp(\hat{\beta}_i)$ , <sup>‡</sup> $(\exp(\hat{\beta}_i) - 1) \times 100$ , <sup>§</sup> Number of eggs /FL (cm)<sup>3</sup> $\times 10^3$

The average return rate over the last five years (2015–2019 releases) was 0.74% in this region, and a 0.1g increase in size-at-release would be expected to increase the return rate to 1.2% (= 0.74 × (1+0.625)) from current levels. A 1°C increase in winter SST in the Gulf of Alaska increased the return rate by 49.9% in the Hokkaido Sea of Japan region. On the contrary, an increase of 100 eggs (relative fecundity) of parent females (age-4) (= egg size reduction) reduced the return rate by 60.6% in the Hokkaido and Honshu Pacific regions, and a 1°C increase in SST at release reduced the return rate by 38.3% in the Honshu Sea of Japan region. An increase in biomass of yellowtail landings of 100 tons predicted an 8.9% reduction in the adult return rate of chum salmon in the Hokkaido Okhotsk region.

The correlation between predicted and observed adult return rates was high in all four coastal regions: Hokkaido Okhotsk (𝑟 = 0.79, *t* = 5.75, 𝑝 = 0.0000), Hokkaido and Honshu Pacific (𝑟 = 0.91, *t* = 9.98, 𝑝𝑝 = 0.0000), Hokkaido Sea of Japan (𝑟 = 0.79, *t* = 5.78, 𝑝 = 0.0003), and Honshu Sea of Japan (𝑟 = 0.81, *t* = 6.17, 𝑝 = 0.0000). Our best models with few (2–4) explanatory variables provided good fits to the four coastal regions and the annual variations in chum return rates, although the 95% prediction intervals are large (Figure 8).

**FIGURE 8.**
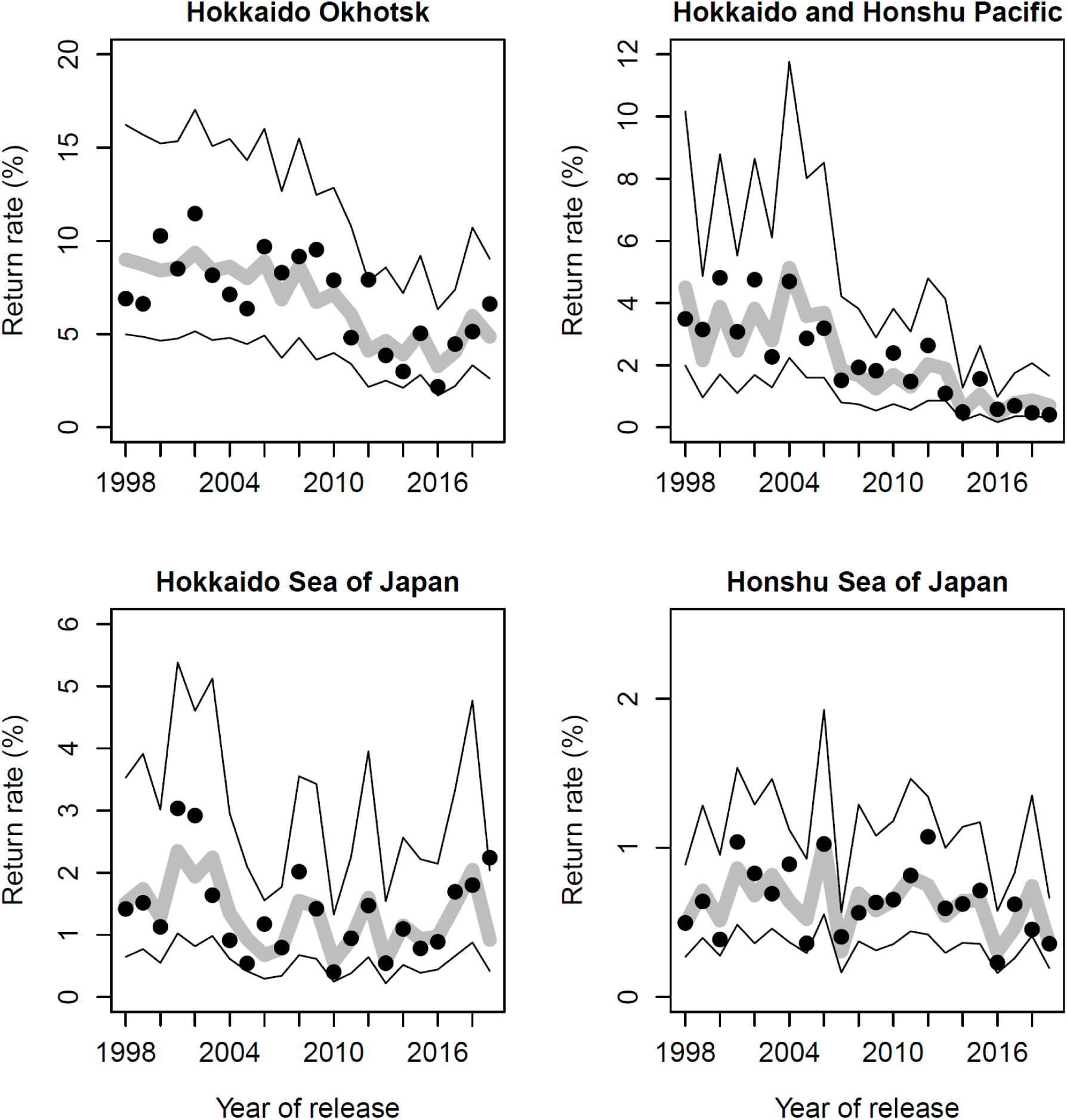
Predicted and observed chum salmon marine survival (adult return rates) by management region in Japan. Predicted return rates (grey line), observed return rates (black dots), and 95% prediction intervals (black lines). Note that scales on return rate axis differ by region. See Figure 1a for management region locations.

## 4| DISCUSSION

We investigated relationships between the marine survival of Japanese hatchery chum salmon and multiple hypothesized stressors during 25 years of climate change. While previous studies of Japanese chum salmon have focused largely on climate change impacts on survival during the early marine (coastal, fry) life history stage, we hypothesized that marine survival is related to climate change effects during all life stages (from eggs to adults). We identified potential climate-driven stressors along known coastal and high-seas migration routes of Japanese chum in the North Pacific Ocean and adjacent seas. Associations between stressors and marine survival varied by coastal region, salmon life stage, and seasonal high-seas distribution area. The stressors exhibiting the largest model effects were fry size-at-release (positive) and parental relative fecundity (negative), and were found only in the Hokkaido and Honshu Pacific where the decline in chum salmon catches was most severe. The negative correlation between relative fecundity and egg size of age-4 fish was significant in this region (*r* = −0.60, *t* = −3.34, *p* = 0.003), indicating that egg size had the largest positive effect. We identified yellowtail, whose distribution was shifting northward, as a new potential marine fish predator of chum salmon fry in the Hokkaido Okhotsk. The model effects of SST were inferred only for fry released in the Hokkaido Sea of Japan and Honshu Sea of Japan regions, and the directional effects of SST were consistent for summer (negative) and winter (positive) in both regions. The SST at release and summer SST in the Sea of Okhotsk and the Bering Sea were negatively associated with marine survival, while winter SST in the western North Pacific and the Gulf of Alaska was positively associated with marine survival. Negative associations between Russian chum and/or Russian pink salmon catches and Japanese chum salmon marine survival were found in all regions, while no associations were identified between US and Canadian chum and pink salmon catches. Moreover, we found a nationwide decline in egg size in Japanese chum salmon over the past two decades.

### 4.1 Adult return rates (marine survival index)

Our adult return rate estimates should be regarded as indices of marine survival because little is known about freshwater mortality of hatchery chum salmon fry after release (e.g., Hasegawa et al., 2021b) or mixtures of regional populations in landed commercial catches in Japan (Nagata et al., 2016). Genetic stock identification showed that 80–98% of the mixed samples of chum salmon caught in September– early October along the western coast of the Sea of Okhotsk originated from Hokkaido rivers, while 53–71% originated from Honshu rivers in late October (Saito et al., 2020). The mixed-stock fishery could bias the return rates in each management region in Hokkaido and Honshu.

It is generally accepted that most chum salmon in Japan are hatchery-reared fish (Hiroi, 1998; Kaeriyama, 1999; Kobayashi, 1980; Iida et al., 2018; Kaeriyama & Sakaguchi, 2023). We assumed all chum salmon returning to Japan were hatchery reared. The total proportion of wild chum salmon in Japan is largely unknown. Field studies provide evidence that natural reproduction of chum salmon occurs in Hokkaido (Miyakoshi et al., 2012) and in the Honshu Sea of Japan (Iida et al., 2018; Iida et al., 2021). Hatchery salmon have a relative reproductive success (RRS, number of offspring a salmon produces compared to other salmon in the population) that is approximately 50% of the RRS of wild fish (Araki et al., 2007; Christie et al., 2014; Shedd et al., 2022), with considerable variation (Kitada et al., 2011). Reduced RRS can be carried over to the next generation (Araki et al., 2009). We expected low freshwater survival from egg to ocean entry of wild chum salmon in natural streams in Japan. Our marine survival indices may be overestimated depending on the magnitude of recruitment of wild fish, which we expected to be very low compared to the number of hatchery-released fish (100–200 million fry per year in each management region, Figure 2). Our regression analysis overestimated the intercept proportional to the magnitude of natural recruitment, but regression coefficients were expected to be robust if natural recruitment and genetic mixture were constant over the period of our analysis.

### 4.2 Regression model assumption

Our linear regression model of the log-transformed return rate assumes that the error is independent and normally distributed. We checked the model residuals to see if the normality assumption of our linear regression model held for our data. The qq plots of the residuals from the best models (Table 2) based on the real scale data showed that most of the studentized residuals from our linear model were within the 95% confidence intervals against the theoretical quantiles of a normal distribution (Figure S17). The result indicates that there are no serious problems with our regression model. This is probably because the catches were not small in number, although the return rates were very small. Return rates are proportions that can also be modeled with a beta distribution with a logit link function. Beta linear regression is a topic for future research. We inferred year effects for logit-transformed return rates with a beta error (Figure S18), which were very similar to those for log-transformed return rates with a normal error (Figure S4), showing that the log-normal model visualizes the year effects of return rates.

### 4.3 Egg size and body size of fry at release

Our finding of a nationwide decline in egg size of age-4 and age-5 Japanese hatchery chum salmon over the past two decades is striking. Egg size is an important fitness trait, and larger eggs have a positive effect on juvenile survival and growth in salmonids (Hutchings, 1991; Einum & Fleming, 1999, 2000). Negative relationships between egg size and fecundity have been reported for Pacific salmon (Beacham & Murray, 1993; Fleming & Gross, 1990). Our regression model coefficients (practical effect size, Table 2) suggested that higher relative fecundity (smaller eggs) could contribute to the reduced survival rate of chum salmon in the Hokkaido and Honshu Pacific region. In fact, the correlation between egg sizes and return rates in this region was significant for age-4 (*r* = 0.76, *t* = 5.17, *p* = 0.0000) and for age-5 (*r* = 0.81, *t* = 6.13, *p* = 0.0000) chum salmon, even though egg size was excluded from the explanatory variables due to multicollinearity (Table S2). Selection in hatcheries could impact fecundity and egg size (Heath et al., 2003), although egg size or fecundity were not found to differ between hatchery and wild fish in other populations (Beacham & Murray, 1993; Quinn et al., 2004; Beacham, 2010). Hasegawa et al. (2021a) showed that the egg size of 5-year-old Japanese chum salmon in 2010 was smaller in more northeasterly locations than in 1994 and 2001, and hypothesized that domestication selection in hatcheries affected by global warming may have caused the temporal shifts in egg size. An alternative process might be maternal effects, whereby environmental variation (e.g., ocean temperature during egg development; maternal diet) experienced by the mother is translated into phenotypic variation in reproductive traits of offspring (e.g., egg size, age at maturation) (reviewed by Mousseau & Fox, 1998; Thorn et al., 2019).

Our regression model coefficients (Table 2) suggested that fry size-at-release and quality of hatchery fish are particularly important for survival at later ocean stages for the Hokkaido and Honshu Pacific region chum salmon. After release, Honshu Pacific region chum fry migrate long distances northward against the Oyashio Current (Figure 1a) to their summer feeding area in the Sea of Okhotsk. On the Sanriku Coast, Honshu Pacific region, where the decline in abundance was most severe (Figure 2), chum salmon fry reared under low food availability (1% body weight) allocated less than half of their energy to growth at high temperatures (14°C) compared to fry reared under high food availability (4% body weight; Iino et al., 2022). In the high food availability group, 32–47% of the energy consumed by the fish should be used for daily growth (Iino et al., 2022). Japanese hatchery chum salmon larger than 2 g (BW) significantly increased long-distance swimming performance and reduced the metabolic cost of swimming against the southward current at 8–12°C SST (Iino et al., 2024). Juvenile chum salmon in western Alaska consumed lower quality prey during warm periods than during cold periods, with the proportion of lower quality prey increasing during warm periods, and the energy density of juvenile chum salmon was lower at warmer SST (Farley et al., 2024).

### 4.4 Predators of fry during coastal migrations

Predation is well-accepted as the proximate cause of salmon fry mortality during their coastal migrations. There are many known fish, bird, and marine mammal predators of Japanese chum salmon (Okado et al., 2020; Hasegawa et al., 2021b; Okado & Hasegawa, 2024). Long-term climate change and marine heatwaves are projected to result in poleward shifts in the distribution of many North Pacific marine species by 2050 (Cheung et al. 2015, Cheung & Frölicher, 2020). Commercial catch data for coastal marine species in Japan was a useful metric for identifying poleward shifts in the distribution of potential piscivorous fish predators of chum salmon fry.

We identified yellowtail as a potential marine fish predator of chum salmon fry. Yellowtail are a subtropical species distributed from Japan and the eastern Korean Peninsula to the Hawaiian Islands (Robins et al., 1991), and are one of the most commercially important marine fish species in Japan. A climate regime shift (warming) during the late 1980s was associated with the northward extension of yellowtail distribution in the Tsushima Warm Current off northwestern Honshu and Hokkaido (Figure 1a), and global warming was projected to further expand the suitable thermal habitat of yellowtail northward (Tian et al., 2012). Since 2014, yellowtail have been caught along the coast of the Sea of Okhotsk in May–December by a setnet fishery targeting adult salmon (September–December), and other marine fish species (May– August) (Hoshino & Fujioka, 2021). The rapid increase in salmon setnet catches of yellowtail led to the identification of this species as a potential indicator of human (fishing) pressure in Japan (Hunsicker et al., 2022). In May 2023, the peak month of hatchery releases of chum salmon fry in the Hokkaido Okhotsk region (Figure S2), schools of yellowtail were often observed on the sea surface near the central Sea of Okhotsk coast (pers. comm., Takashi Ichikawa). Adult yellowtail are well-known to feed on small pelagic fish and invertebrates (Robins et al., 1991), but stomach contents surveys are necessary to confirm yellowtail predation on chum salmon fry.

Our regression model results indicated yellowtail catches are an important covariate of fry chum survival in the year of release only in the Hokkaido Okhotsk region, although yellowtail catches were higher in the more southerly management regions such as Hokkaido Sea of Japan and Honshu Pacific (Figure 5b). Yellowtail spawning grounds cover a wide area from the East China Sea to western Kyushu Island, and the main spawning season is from February to June (Tian et al., 2012). After spawning, adult yellowtail migrate north for feeding. Yellowtail fry also migrate north on drifting seaweed using the Tsushima Warm Current in the Sea of Japan and the Kuroshio Current off the Pacific coast of Honshu (Figure 1a). Yellowtail (age 0–3 years and older) are caught in Hokkaido from May to August (Hoshino & Fujioka, 2021), while hatchery chum fry are released mainly in March-April in the Hokkaido Sea of Japan region (Figure S2). On the Pacific coast of Honshu, yellowtail (age unknown) are caught in commercial fisheries from late July to mid-September (Iwate Prefecture, unpublished figure) and from June to December (Miyagi Prefectural Government, 2022), while hatchery chum fry were released mainly in February–April (Figure S2). The earlier timing of chum fry release compared to the timing of the yellowtail migration may be a reason for the exclusion of yellowtail catches from the explanatory variables in these regions.

Annual estimates of stock abundance (biomass) or relative abundance (catch per unit effort) would be more appropriate covariates than annual catches when considering many potential predator species (e.g., Pauly et al., 2013; Thorson et al., 2020; Punt, 2023), but were not available for the period of our study. The use of stock assessment data is a promising approach for developing new indices of climate-driven predator exposure, especially if predator diet information is available. A similar approach for high-seas migration areas may be possible through future scientific collaborations among international treaty organizations (PICES-NPFC, 2019).

### 4.5 Inter- and intra-specific competition during high-seas migrations

Indirect evidence of competition between Russian pink salmon and many other marine species, including chum salmon, is substantial (see review by Ruggerone et al., 2023). Our results suggest that Russian pink and chum salmon compete for food, space, or other resources with Japanese chum salmon (ages 2–3) in the central Bering Sea in summer and the Gulf of Alaska in winter, while competition with immature (age-1) chum salmon could be weak or negligible in the Sea of Okhotsk and western North Pacific (Naydenko & Temnykh, 2016; Naydenko et al., 2016; Bugaev & Gerlits, 2023). Our analysis found no relationship between Japanese chum salmon return rates and North American pink and chum salmon catches. This may reflect inter- and intra-specific differences in the ocean distribution of Japanese chum salmon and North American pink and chum salmon. For example, limited data from salmon research vessel surveys in winter indicate Japanese chum salmon are distributed farther to the south in the Gulf of Alaska and northeastern North Pacific than North American chum salmon (reviewed by Myers et al., 2016; Urawa et al., 2016). Past research indicated negative intraspecific relationships between the abundance of Japanese hatchery chum salmon and the abundance and annular scale growth of local and regional stock groups of North American chum salmon (Ruggerone et al., 2011; Agler et al., 2013; Frost et al., 2021). All our best regional models included North American chum salmon as an explanatory variable with VIF < 3 (Table S2), but were excluded by our model selection procedure. Our regression analysis (Table 2) detected a negative association between return rates of Hokkaido Sea of Japan chum and Russia chum catch (*r* = −0.44, *t* = −2.22, *p* = 0.038), while correlations between regional return rates and North American chum catch were too weak to detect North American chum salmon as a competitor.

A reanalysis of coastal and high-seas salmon research survey data (1953–2022) confirmed the results of earlier studies showing the overlap in distributions of chum, pink, and sockeye salmon throughout the North Pacific (Langan et al., 2024). A meta-analysis of salmon trophic ecology data showed similar diet composition of chum, pink, and sockeye salmon throughout the North Pacific, with a diet overlap of 31.8% between chum and pink and 30.9% between chum and sockeye, and diet overlap was highest between pink and sockeye salmon at 46.6% (Graham et al., 2021). The estimated proportion of Asian and North American hatchery chum salmon was 60% of the total abundance of maturing chum salmon during 1990–2015 (Ruggerone & Irvine, 2018). The substantial decline in abundance of Japanese chum salmon after the mid-2000s may have reduced inter- and intra-specific competition for prey resources with Russian chum and pink salmon. The lack of an association between Russian pink salmon catches (*t*+1) and survival of age-1 Japanese chum salmon might also indicate improved growth and survival in the Sea of Okhotsk. However, even when catches of Japanese chum salmon were low, their survival (age-2 and age-3 fish) was negatively correlated with Russian chum and pink salmon catches, suggesting that Russian chum and pink salmon have a competitive advantage over Japanese chum salmon.

Directional swimming to preferred thermal habitat seems to play an important role in the seasonal northward migrations of salmon (Azumaya & Ishida, 2004). Russian pink and chum salmon may have a competitive advantage simply because they do not have as far to swim from their overwintering habitat (central North Pacific and Aleutian Islands) as Japanese chum salmon (Gulf of Alaska) to their shared optimal summer feeding habitat in the central Bering Sea. If Russian salmon are first to arrive in the central Bering Sea in summer, they might occupy most of the optimal thermal habitat and reduce the amount of available preferred prey, causing a trophic cascade (Batten et al., 2018). Models project that the optimal thermal habitat of chum and pink salmon will continue to contract and shift northwestward under future climate-change scenarios (e.g., Abdul-Aziz et al., 2011, also see Figures 4, S7), and we expect Russian salmon to continue to have a competitive advantage over Japanese chum salmon.

We hypothesize that inferior athletic performance in a warming climate may be related to the long-term decline in the marine survival of Japanese hatchery chum salmon. Wild-born chum salmon fry were too agile to be netted before 1945 in a river on Etrofu Island where Japan operated salmon hatcheries (Yagisawa, 1970). Currently, piscivorous salmonids chase schools of hatchery chum salmon fry, and small fry are easily predated due to their slow swimming speed to escape predators (Hasegawa et al., 2021b). Water temperature and oxygen are important regulators of physiological processes in fish (Powers et al., 1991; Somero, 2022). Swim tunnel experiments show that temperature and body size influence swimming performance in Japanese hatchery chum fry (Iino et al., 2024). Swimming performance has been least studied in adult chum salmon and Chinook salmon (reviewed by Kraskura et al., 2024).

### 4.6 Ocean conditions in coastal and high-seas habitats

Ranges of chum salmon thermal habitat used in our analysis were narrower (thermal limits, 4– 13°C) than previously published estimates, based largely on historical high-seas salmon survey data collected before the winter 1976–1977 climate regime shift (preferred spring-summer SST range, 1.3–12.6°C; Langan et al., 2024). The pre-regime shift period (1948–1975) was characterized by low ocean productivity, cold winter SSTs in the Gulf of Alaska, and a low abundance of salmon populations that overwinter in the Gulf of Alaska, including Japanese chum salmon. During periods of low ocean productivity, optimal temperatures for salmon growth decline (Beauchamp, 2009).

Marine heatwaves and long-term climate change are projected to result in poleward shifts in the distribution of many North Pacific pelagic fish species by 2050 (Cheung et al. 2015, Cheung & Frölicher, 2020). Our maps of Japanese chum salmon thermal habitat indicated a summer northward shift and contraction since 2010; winter habitat remained relatively stable with a slight northeastward shift in 2017 and 2018 (unpublished figure, S. Kitada). The frequency of annual marine heatwaves increased in the western and central Bering Sea, and heatwaves in the eastern Bering were longer and more intense in the past decade (2010–2019) (Carvalho et al., 2021). Marine heatwaves in the Gulf of Alaska (2013–2016, 2018–2020) were accompanied by increased acidity and low oxygen, resulting in cascading ecosystem effects (Hauri et al., 2024). Our analysis showed increases in SST in the central Bering Sea increased after 2010 and SST in the Gulf of Alaska in 2014–2015 and 2019–2020 (Figure S11b) that are consistent with the occurrence of marine heat waves.

Japanese chum salmon from all regions would experience similar environmental effects from common coastal and high-seas habitat conditions, but the effects of SST were inferred only for fry released in the Hokkaido Sea of Japan and Honshu Sea of Japan regions. The Sea of Japan is a global hotspot for long-term declines in marine fisheries productivity (Free et al., 2019). After a brief (1998–1999) natural cooling event, anthropogenic climate change became the principal driver of extreme warming events in the Sea of Japan, particularly in southern regions (Hayashi et al., 2022). Our regression analysis found that SST at release was negatively associated with return rates in the Honshu Sea of Japan. Summer SST (July) on the Hokkaido Okhotsk coast was also negatively associated with return rates for fry released in the Sea of Japan regions. Summer SST (July) on the Hokkaido Okhotsk coast (range, 10.8–14.2°C) was higher than published temperatures for optimal growth and feeding (8–12°C) and available habitat for feeding (5–13°C) of Japanese chum salmon (Kaeriyama et al., 2012; Urawa et al., 2018). Hatchery chum salmon fry were released mainly in February–March along the Honshu Sea of Japan coast and March–April along the Hokkaido Sea of Japan coast (Figure S2). After release, chum salmon juveniles migrate in the direction of the northbound Tsushima Warm Current to the Sea of Okhotsk (Figure 1a). The negative association of survival with July SST may be related to lengthy exposure to warm coastal SSTs in the Sea of Japan and delayed arrival at their nursery area in the Sea of Okhotsk. Juvenile chum salmon were abundant in the coastal Okhotsk Sea from May to June when SST was between 8 and 13°C, and disappeared from coastal waters after late June when SST exceeded 13°C (Nagata et al., 2007). Hokkaido Okhotsk is the most successful management region (Figure 2), and the peak release time is in May (SST, 4.7–6.2°C). Late arrival of juveniles released from the Honshu and Hokkaido Sea of Japan regions to the Sea of Okhotsk may reduce survival during the first summer. The results suggest that release timing is important to avoid mortality at release and summer mortality on the Hokkaido Okhotsk coast.

In high-seas habitats, summer SST (August–September) in the Bering Sea (range, 8.3–11.4°C) was within the published optimal growth and feeding range of Japanese chum salmon (8–12°C; Kaeriyama et al., 2012; Urawa et al., 2018). The negative effect of summer SST in the Bering Sea was found only for fry released in the Hokkaido Sea of Japan. In this region, egg size was the smallest in Japan (Figure S15). These results suggest a hypothesis that the upper thermal tolerance is lower for hatchery salmon produced in this region. On the contrary, the model effects of winter SST in the western North Pacific and in the Gulf of Alaska were consistently positive for fry released in the Sea of Japan regions. Winter SST (January–April) increased after 2010 in the western North Pacific (range, 4.6–6.1°C) and the Gulf of Alaska (range, 4.6–7.2°C) (Figure S11b); these were similar or slightly higher than the published overwintering temperature range (4–6°C; Kaeriyama et al., 2012; Urawa et al., 2018). This result supports the hypothesis that Japanese chum salmon in the Sea of Japan regions affected by the Tsushima Warm Current are better able to survive increasing winter SST (Figure S11b).

In 2022, the number of chum salmon returning to Japan unexpectedly increased in the Hokkaido Sea of Japan and the Hokkaido Okhotsk management regions (Figure 2). In the Chitose River in the Hokkaido Sea of Japan, where Japan’s oldest hatchery operates, 587,000 fish returned, surpassing the historical record of 551,000 in 1995 (Chitose Aquarium, 2023). The sudden increase in 2022 surprised fishermen, hatchery managers, local people, and biologists. However, returning chum numbers declined again in all geographic regions in 2023, reaching historic lows (a total of about 18 million in Japan) in 2024. In the Hokkaido Sea of Japan and Okhotsk regions, the peak in adult returns in 2022 (Figure 2) was mainly caused by an increase in age-4 fish released as fry in 2019 (Figure S19), indicating better survival of the 2019 year class. Thermal habitat (8–10°C) within the published optimal thermal habitat range for growth and feeding (8–12°C) of Japanese chum salmon was widely distributed in the Sea of Okhotsk in July 2019 and in the central Bering Sea in September 2020 (Figure S20), where age-1 Japanese chum salmon are distributed during their second summer growing season. Expansion of optimal thermal habitat during the first and second summers likely improved the survival of immature chum salmon in 2020 and 2021, and may have resulted in the anomalous peak returns of age-4 fish in 2022. In contrast, total return rates decreased in other regions (Honshu Sea of Japan and the Pacific regions; Figure S19). The results suggest that coastal mortality of fry released from these regions was higher and substantial.

### 4.7 Management implications

The largest positive model effect identified by our regression analysis was fry size-at-release and egg size (relative fecundity, negative), suggesting that hatchery fry quality (size-at-release) and release timing are particularly important for survival in later ocean stages. Hatchery experiments to test appropriate fry size and time of release and nearshore monitoring of SSTs might provide an adaptive approach for improving adult returns of Japanese chum salmon (Saito & Nagasawa, 2009; Nagata et al., 2016; Iida et al., 2021; Saito, 2022; Iida, 2024; Honda et al., 2025). Adjustment of release timing to allow fry to arrive in the Sea of Okhotsk in May or early June or both might improve nearshore survival. Experiments to assess fry quality, metabolism, and swimming performance of Japanese chum salmon (e.g., Iino et al. 2022, 2024; Abe et al., 2024) should provide information useful for improving hatchery fry quality.

International comparisons of the swimming performance of hatchery and wild chum salmon may provide crucial information for Japanese chum salmon fitness. Moreover, our analysis found a nationwide decline in egg size over the past two decades. The mechanisms of egg size reduction in Japanese chum salmon are unknown and an important topic for further study. A nationwide evaluation of current hatchery techniques including parent selection, egg incubation, and fry rearing and release strategies would be useful for understanding the underlying mechanisms of the decline of Japanese chum salmon abundance over the past two decades.

Investigations of many other potentially important stressors beyond the scope of our study remain to be addressed. For example, relationships between the marine survival of salmon and climate-driven changes in marine fisheries operating along high-seas salmon migration routes and within the 200-mile EEZs of other nations. In Alaska, genetic analyses show that Asian (Russia and Japan) chum salmon populations dominate early (June) catches by salmon fisheries targeting sockeye salmon along the south side of the Alaska Peninsula in early (June), when chum salmon are migrating from winter (North Pacific) to summer (Bering Sea) feeding grounds (Dann et al., 2023). Asian stocks of chum salmon also dominate prohibited species bycatch of salmon in the US walleye pollock (*Gadus chalcogrammus*) fishery in the eastern Bering Sea in early summer (Barry et al., 2022). Bycatches of Asian chum salmon (ocean-age-3 and age-4; unknown maturity stage) by both fisheries in 2021 were unexpectedly high (Barry et al., 2022; Dann et al., 2023). The Alaska bycatch of Asian chum salmon preceded the spike in adult returns to Japan in 2022. The chum salmon bycatch in the Bering Sea decreased substantially in 2022 and 2023 (NPFMC, 2025), coinciding with the decrease in adult returns to Japan in 2023 and 2024. Seeking predictive (vs. retrospective) indices would benefit preseason forecasting of adult salmon returns. In the absence of quantitative marine stock assessment surveys of salmon, potentially predictive indices might be used in a qualitative “stoplight” approach to pre-season forecasting (Caddy, 2002; NOAA, 2024). This method requires an adaptive approach because relationships between salmon survival and index variables change over time (non-stationarity; Litzow et al., 2018; Ohlberger et al., 2022).

## 5| CONCLUSIONS

Our results from the world’s largest long-term salmon hatchery program indicate that maternal traits (egg size and fecundity), hatchery fry quality, fry body size-at-release, and release timing are particularly important for survival at later ocean stages. Egg size and parental fecundity are key demographic parameters that contribute to the fitness and productivity of salmon populations, and the decadal-scale changes observed herein could result in significant ecosystem, fishery, and societal losses. Investigation of the mechanisms of egg size reduction is one of the most important topics for further study in Japanese chum salmon. Our study highlights the need for an experimental approach to hatchery practices, including monitoring and analyses with updated information, leading to effective management decisions and policies for future sustainability and conservation of salmon resources.

## AUTHOR CONTRIBUTIONS

Shuichi Kitada: Conceptualization (lead); data curation (lead); formal analysis (lead); funding acquisition (lead); investigation (lead); methodology (equal); software (lead); validation (equal); visualization (lead); writing–original draft (lead); writing–review and editing (equal). Katherine W. Myers: Conceptualization (equal); writing–review and editing (lead). Hirohisa Kishino: Conceptualization (equal); formal analysis (support); funding acquisition (equal); methodology (lead); validation (lead); visualization (support); writing–review and editing (equal). All authors approved the final version of the manuscript.

## ACKNOWLEDGMENTS

We sincerely acknowledge the thousands of biologists, fishery and hatchery managers, and fishers who contributed to the collection of data analyzed in our manuscript. The efforts of APG, FRA, HSEA, JMA, MAFF, NOAA, and NPAFC to create public databases made our study possible. We thank Brian Riddell and the reviewers for their insightful and constructive comments, which significantly improved the earlier version of the manuscript. Takashi Ichikawa provided information on chum salmon and yellowtail. This study was supported by Japan Society for the Promotion of Science Grants-in-Aid for Scientific Research KAKENHI No. 18K0578116 for SK and No. 22K11950 for HK.

## CONFLICT OF INTEREST STATEMENT

The authors declare that no conflict of interest exists.

## DATA AVAILABILITY STATEMENT

The data generated or analyzed in this study are fully described in the published article and its Supplementary Information. Analysis scripts and data will be available at https://doi.org/10.5281/zenodo.14868156 on acceptance.

## Notes

https://doi.org/10.5281/zenodo.14868156

### Competing Interest Statement

The authors have declared no competing interest.

### Summary of Updates

All sections thoroughly revised to improve clarity of statistical methods and results; model assumptions tested; Table 3 newly created to report practical effect sizes; Figures 5(a) and 7 revised; Figures S3, S4, S13, S15, S17, S18 newly created; analysis of age at maturity and body size removed; Discussion and Conclusions extensively revised.

