## Supplementary Information for "Hatcheries to high seas: climate change connections to salmon marine survival"

#### Appendix S1

### Supplementary Information for:

### Hatcheries to high seas: climate change connections to salmon marine survival

Shuichi Kitada, Katherine W. Myers, and Hirohisa Kishino

This PDF file includes:

Tables S1–S4

Figures S1–S20

The R scripts and Data S1–S7

#### Tables S1–S4

**Table S1** Results of the generalized additive models (GAMs) fitted to explanatory variables across years (1998-2019 releases of hatchery chum salmon fry) by management region in Japan. Bold indicates significant fitting of nonlinear and/or linear year effect (α=0.05). Sea surface temperature (SST). See Figure 1a for management region locations.

| Management region | log (Return rate) | | SST at release | | Fry size at release | | Egg size (age-4) | | Egg size (age-5) | |
|---|---|---|---|---|---|---|---|---|---|---|
| | $p$ | $R^2$ | $p$ | $R^2$ | $p$ | $R^2$ | $p$ | $R^2$ | $p$ | $R^2$ |
| Hokkaido Okhotsk | **0.0015** | **0.64** | 0.5040 | 0.02 | **0.0000** | **0.87** | **0.0148** | **0.54** | **0.0016** | **0.51** |
| Hokkaido Nemuro | **0.0000** | **0.81** | **0.0055** | **0.29** | **0.0001** | **0.79** | **0.0008** | **0.53** | **0.0001** | **0.63** |
| Hokkaido Pacific East | **0.0000** | **0.55** | 0.1330 | 0.17 | 0.1130 | 0.25 | **0.0387** | **0.29** | 0.0538 | 0.26 |
| Hokkaido Pacific West | **0.0001** | **0.63** | **0.0111** | 0.40 | 0.8600 | -0.05 | **0.0034** | **0.35** | **0.0063** | **0.39** |
| Hokkaido Sea of Japan | 0.1730 | 0.19 | 0.6630 | -0.04 | **0.0017** | **0.70** | **0.0011** | **0.48** | **0.0000** | **0.73** |
| Honshu Pacific | **0.0000** | **0.90** | **0.0007** | **0.59** | **0.0001** | **0.82** | **0.0028** | **0.34** | **0.0077** | **0.46** |
| Honshu Sea of Japan | 0.2930 | 0.08 | 0.2860 | 0.11 | **0.0009** | **0.49** | 0.3280 | 0.08 | 0.1480 | 0.06 |

**Table S2** Model selection for subset models of hypothesized explanatory variables on Japanese chum salmon adult return rate (marine survival) by management region in Japan. Bold indicates the best model in each management region. See Figure 1a for region locations. See Table S3 for definitions of abbreviations of excluded variables. Number of parameters to be estimated (No. para.); Variance inflation factor (VIF); Akaike Information Criterion, small sample correction (AICc; Sugiura, 1978)

| Region | Subset model | No. Para. | Excluded variables with multicollinearity (VIF > 3) | Using all explanatory variables (VIF < 3) | | | | Best models after variable selection | | |
|---|---|---|---|---|---|---|---|---|---|---|
| | | | | VIFmax | No. para. | AICc | Adjusted $R^2$ | No. para. | AICc | Adjusted $R^2$ |
| Hokkaido Okhotsk | Hatchery carryovers | 3 | † Egg4 | 1.30 | 2 | 32.68 | -0.08 | 0 | 27.55 | – |
| | Ocean conditions | 5 | | 1.96 | 5 | 36.89 | 0.09 | 1 | 26.17 | 0.13 |
| | **Predators & competitors** | 9 | RUSchum, AMpink2, Ampink | 1.96 | 6 | 27.76 | 0.49 | 2 | **11.79** | **0.58** |
| | Hatchery carryovers + Ocean condition | 8 | Egg4 | 2.05 | 7 | 46.95 | 0.00 | 1 | 26.17 | 0.13 |
| | Hatchery carryovers + Predators & competitors | 12 | Egg4, RUSchum, AMpink2, Ampink | 2.25 | 8 | 36.74 | 0.51 | 2 | **11.79** | **0.58** |
| | Ocean condition + Predators & competitors | 14 | GoaJanApr, Yellowtail, RUSchum, AMpink, AMpink2 | 2.86 | 9 | 56.37 | 0.11 | 1 | 25.86 | 0.14 |
| | Full model | 17 | Egg4, GoaJanApr, RUSchum, RUSpink, AMpink, AMpink2 | 2.87 | 11 | 62.14 | 0.52 | 2 | **11.79** | **0.58** |
| Hokkaido and Honshu Pacific | Hatchery carryovers | 3 | | 2.67 | 3 | 39.79 | 0.62 | 2 | 39.20 | 0.59 |
| | Ocean conditions | 5 | | 2.00 | 5 | 63.39 | 0.14 | 1 | 52.64 | 0.18 |
| | Predators & competitors | 9 | RUSchum, AMpink, AMpink2 | 2.04 | 6 | 52.30 | 0.56 | 2 | 38.17 | 0.61 |
| | Hatchery carryovers + Ocean condition | 8 | Egg4 | 2.42 | 7 | 54.41 | 0.60 | 3 | 37.97 | 0.65 |
| | **Hatchery carryovers + Predators & competitors** | 12 | Egg4, RUSchum, AMpink, AMpink2 | 2.75 | 8 | 46.02 | 0.79 | 4 | **28.99** | **0.79** |
| | Ocean condition + Predators & competitors | 14 | GoaJanApr, Yellowtail, RUSchum, RUSpink, AMpink, AMpink2 | 1.48 | 8 | 68.67 | 0.41 | 2 | 44.51 | 0.48 |
| | Full model | 17 | Egg4, GoaJanApr, Yellowtail, RUSchum, RUSpink, AMpink, AMpink2 | 2.17 | 10 | 59.58 | 0.80 | 4 | **28.99** | **0.79** |

**Table S2 (continued)**

| Region | Subset model | No. Para. | Excluded variables with multicollinearity (VIF > 3) | Using all explanatory variables (VIF < 3) | | | | Best models after variable selection | | |
|---|---|---|---|---|---|---|---|---|---|---|
| | | | | VIFmax | No. para. | AICc | Adjusted $R^2$ | No. para. | AICc | Adjusted $R^2$ |
| Hokkaido Sea of Japan | Hatchery carryovers | 3 | | 1.36 | 3 | 46.59 | -0.14 | 0 | 38.07 | – |
| | Ocean conditions | 5 | | 1.94 | 5 | 35.77 | 0.46 | 3 | 30.67 | 0.45 |
| | Predators & competitors | 9 | RUSchum, AMpink, AMpink2 | 2.01 | 5 | 55.80 | -0.13 | 0 | 38.07 | – |
| | Hatchery carryovers + Ocean condition | 8 | BerAugSep | 1.62 | 7 | 50.27 | 0.28 | 2 | 33.55 | 0.31 |
| | Hatchery carryovers + Predators & competitors | 12 | Egg4, RUSchum, AMpink, AMpink2 | 2.70 | 8 | 66.76 | -0.19 | 0 | 38.07 | – |
| | **Ocean condition + Predators & competitors** | 14 | Yellowtail, RUSpink, Ampink, AMpink2 | 2.67 | 10 | 64.06 | 0.47 | 4 | **29.42** | **0.54** |
| | Full model | 17 | BerAugSep, Yellowtail, RUSpink, AMpink, AMpink2 | 2.55 | 12 | 99.22 | 0.15 | 3 | 32.24 | 0.41 |
| Honshu Sea of Japan | Hatchery carryovers | 3 | | 1.13 | 3 | 32.25 | -0.07 | 0 | 25.09 | – |
| | Ocean conditions | 5 | | 1.99 | 5 | 22.92 | 0.46 | 2 | 14.85 | 0.46 |
| | Predators and competitors | 9 | RUSchum, AMpink, AMpink2 | 2.08 | 6 | 37.60 | 0.11 | 1 | 19.12 | 0.29 |
| | Hatchery carryovers + Ocean condition | 8 | | 2.58 | 8 | 26.80 | 0.65 | 2 | 14.85 | 0.46 |
| | Hatchery carryovers + Predators & competitors | 12 | AMpink, AMpink2 | 2.69 | 10 | 59.13 | 0.23 | 3 | 19.11 | 0.41 |
| | **Ocean condition + Predators & competitors** | 14 | Yellowtail, RUSpink, AMpink, AMpink2 | 2.98 | 10 | 47.41 | 0.55 | 4 | **14.59** | **0.57** |
| | Full model | 17 | GoaJanApr, Yellowtail, RUSchum, RUSpink, AMpink, AMpink2 | 2.99 | 11 | 54.51 | 0.62 | 2 | **14.59** | **0.57** |

† Egg size (Egg) and relative fecundity (RF) of age 4 and age 5 fish were examined, but these variables were not selected in all cases except for Hokkaido and Honshu Pacific (Table 2), where age 4 fish had a smaller AICc (30.0) and larger $R^2$ (0.79) than age–5 fish (AICc =38.2 and $R^2$=0.61). Therefore, we used Egg4 and RF4 in the analysis of the subset models.

**Table S3** Abbreviations for explanatory variables in Table S1. *t* = hatchery release year of chum salmon fry. SST = sea surface temperature.

| Subset models | Explanatory variables | Abbreviation |
|---|---|---|
| Hatchery carryovers | Relative fecundity in year *t* -1 (Age 4 and Age 5) | RF4 and RF5 |
| | Egg size in year *t* (Age 4 and Age 5) | Egg4 and Egg5 |
| | Size at release in year *t* | SAR |
| Ocean conditions | SST at release in year *t* | SSTrel |
| | Sea of Okhotsk summer SST in year *t* | OkhJul |
| | North Pacific Winter SST in year *t*+1 | NpoJanApr |
| | Bering Sea summer SST in year *t*+1 | BerAugSep |
| | Gulf of Alaska winter SST in year *t*+2 | GoaJanApr |
| Predators and competitors | Yellowtail catch of in year *t* | Yellowtail |
| | Russian chum catch in year *t*+3 | RUSchum |
| | Russian pink catch in year *t*+1 | RUSpink |
| | Russian pink catch in year *t*+2 | RUSpink2 |
| | Russian pink catch in year *t*+3 | RUSpink3 |
| | US and Canadian chum catch in year *t*+3 | AMchum |
| | US and Canadian pink catch in year *t*+1 | AMpink |
| | US and Canadian pink catch in year *t*+2 | AMpink2 |
| | US and Canadian pink catch in year *t*+3 | AMpink3 |

**Table S4** Model averaging of Japanese chum salmon survival in four coastal regions in Japan and regression coefficients with their $p$-values in parentheses. Model averaging was performed on subset models with ΔAICc values less than 2. Bold indicates significant relationship ($\alpha = 0.05$). $t$ = hatchery release year of chum salmon fry. See Figure 1a for management region locations.

| Subset model | Explanatory variable (year) | Management region | | | |
|---|---|---|---|---|---|
| | | Hokkaido Okhotsk | Hokkaido & Honshu Pacific † | Hokkaido Sea of Japan | Honshu Sea of Japan |
| Hatchery carryovers | Relative fecundity † ‡ (*t*-1) | – | **–0.931 (0.0009)** | – | – |
| | Egg size † (*t*-1) | – | – | – | – |
| | Size at release (*t*) | – | **0.486 (0.0009)** | – | – |
| Ocean conditions | SST at release (*t*) | – | – | | **-0.517 (0.0025)** |
| | Okhotsk Jul SST (*t*) | – | – | **-0.326 (0.0194)** | -0.139 (0.0633) |
| | North Pacific Jan-Apr SST (*t*+1) | – | – | 0.292 (0.1263) | 0.238 (0.1077) |
| | Bering Sea Aug-Sep SST (*t*+1) | – | – | **-0.387 (0.0158)** | – |
| | Gulf of Alaska Jan-Apr SST (*t*+2) | – | – | **0.391 (0.0045)** | – |
| Predators and competitors | Yellowtail catch in year (*t*) | **-0.095 (0.0000)** | – | – | – |
| | Russia chum catch (*t*+3) | – | – | **-0.150 (0.0487)** | – |
| | Russia pink catch (*t*+1) | – | – | – | – |
| | Russia pink catch (*t*+2) | -0.127 (0.0566) | **-0.311 (0.0025)** | – | -0.135 (0.0895) |
| | Russia pink catch (*t*+3) | – | **-0.225 (0.0138)** | – | -0.090 (0.2268) |
| | USA & Canada chum catch (*t*+3) | | | -0.259 (0.1903) | |
| | USA & Canada pink catch (*t*+1) | – | – | – | – |
| | USA & Canada pink catch (*t*+2) | – | – | – | – |
| | USA & Canada pink catch (*t*+3) | – | – | – | – |

The three subset models were fitted to log return rates for 1998-2019 releases.

† No other model had ΔAICc values below 2, and the result of the best model was presented.

#### Figures S1–S20

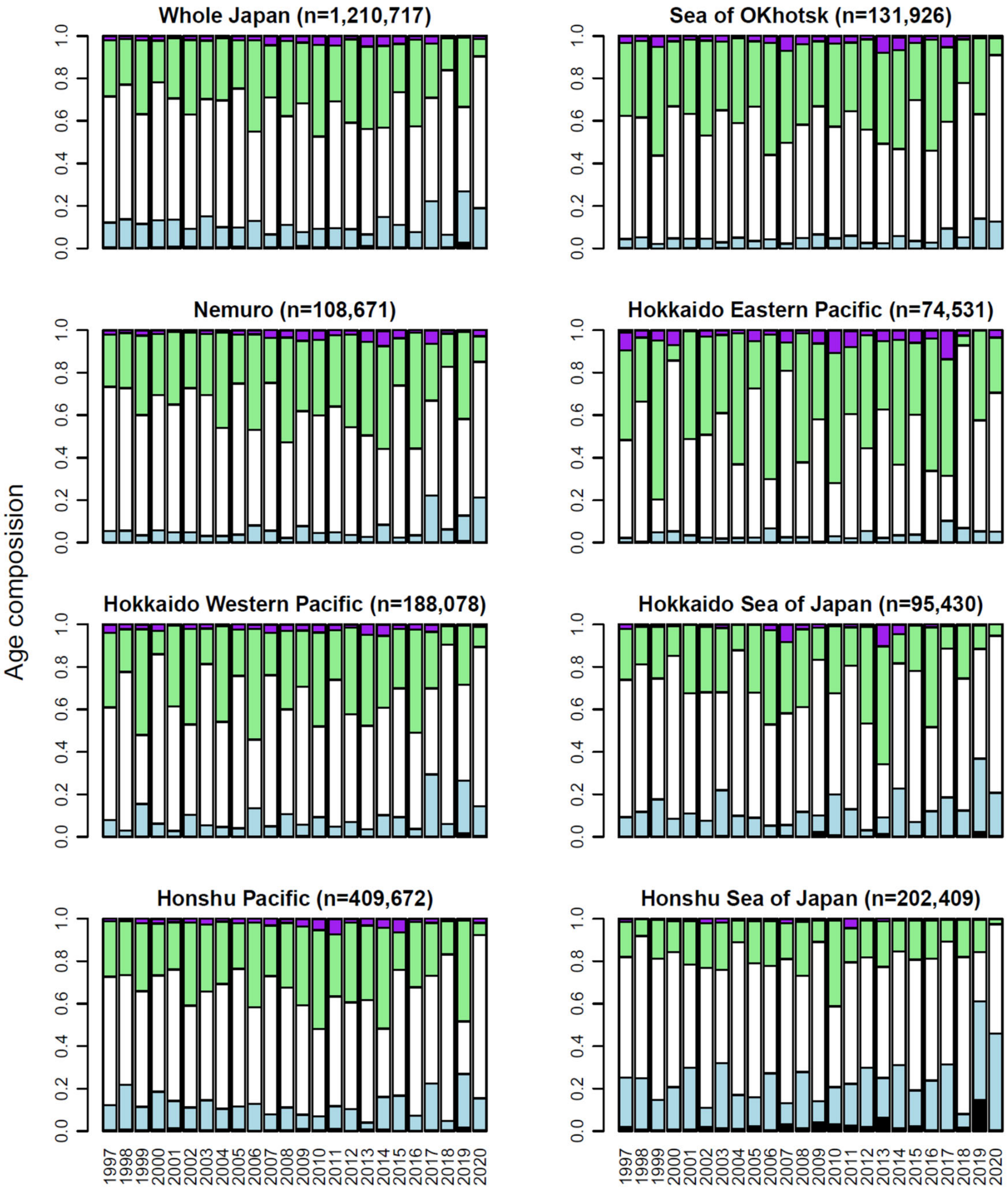


**Figure S1** Age composition of adult chum salmon returns by management area in Japan, 1997-2020. Numbers in parentheses indicate sample size. Black: age-2, light blue: age-3, white: age-4, green: age-5, purple: age-6 and older. See Figure 1a for management region locations.

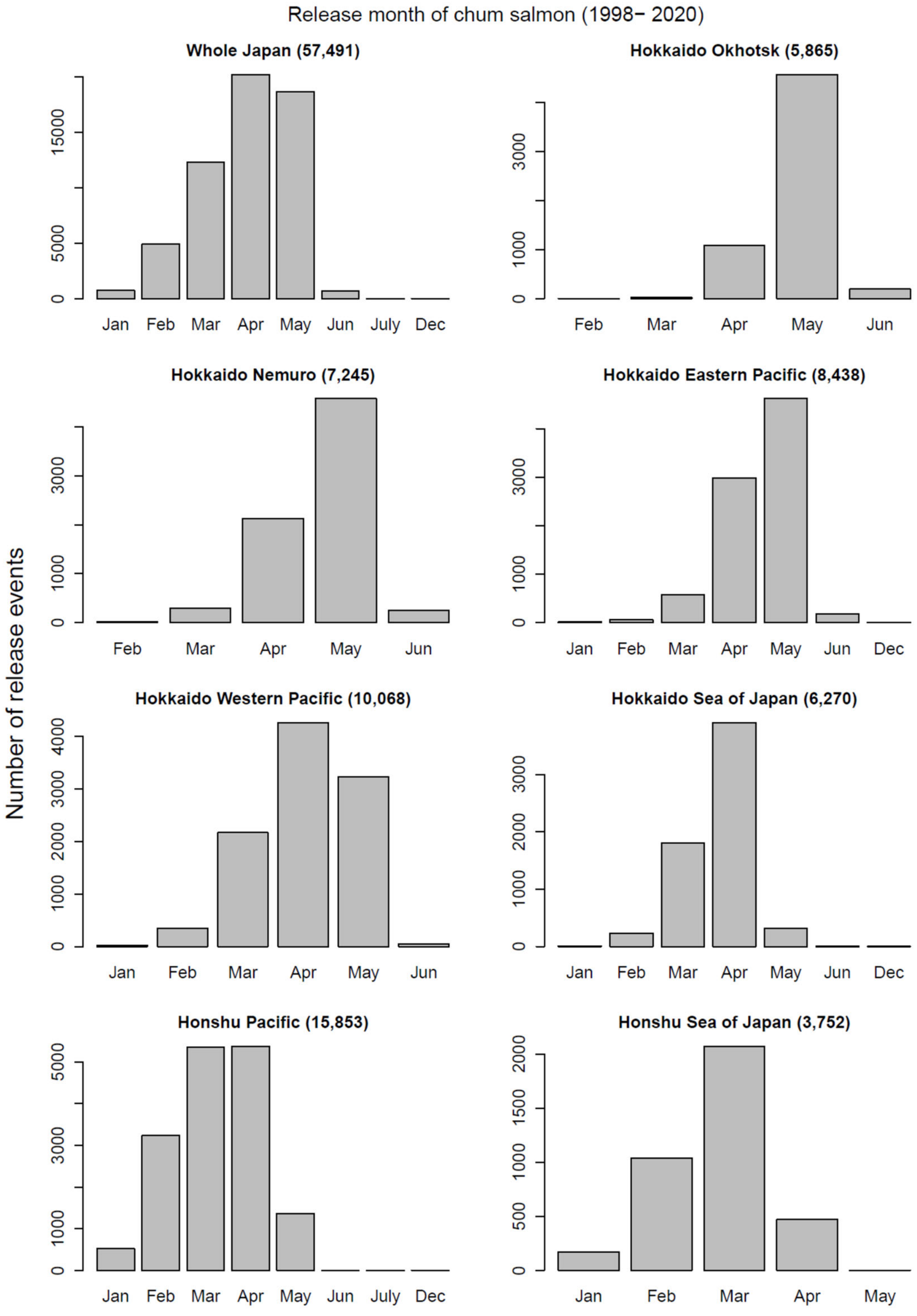


**Figure S2** Month of hatchery chum salmon fry release by management area in Japan, 1998-2020. Numbers in parentheses indicate number of release events. See Figure 1a for management region locations.

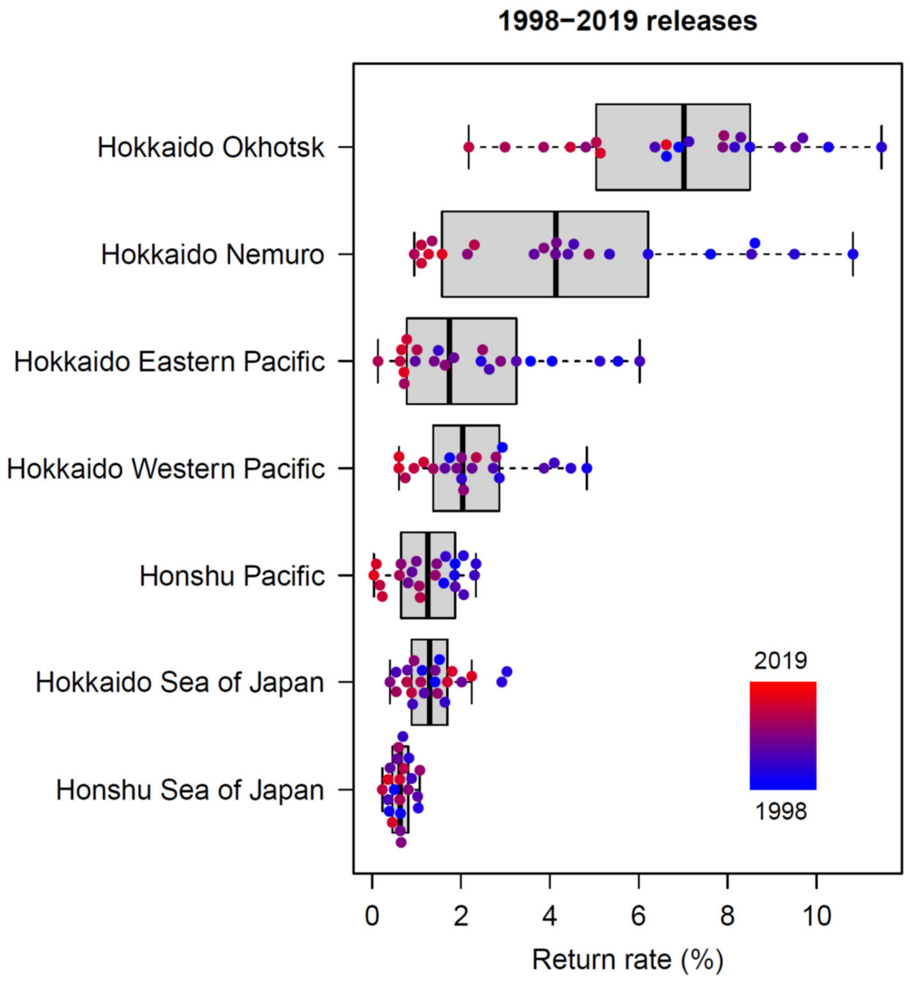


**Figure S3** Chum salmon marine survival (adult return rate) varies by management region in Japan. Adult return rates calculated for age 2–5 adult returns to seven management regions in Japan for fry released in 1998–2019. See Figure 1a for management region locations.

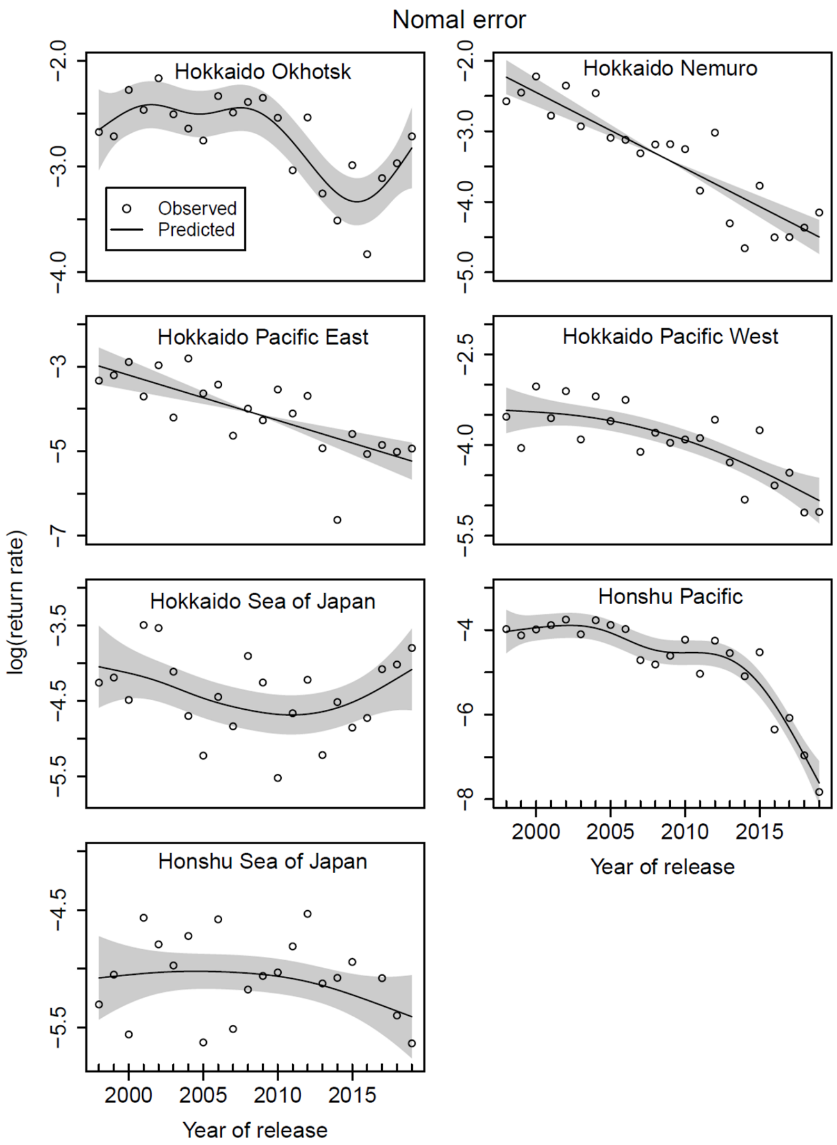


**Figure S4** Log-transformed chum salmon adult return rates by management region in Japan, 1998–2019, as shown by the nonlinear and/or linear year effect from generalized additive models (GAMs) (Table S1). Shaded areas show 2×standard errors. See Figure 1a for management region locations.

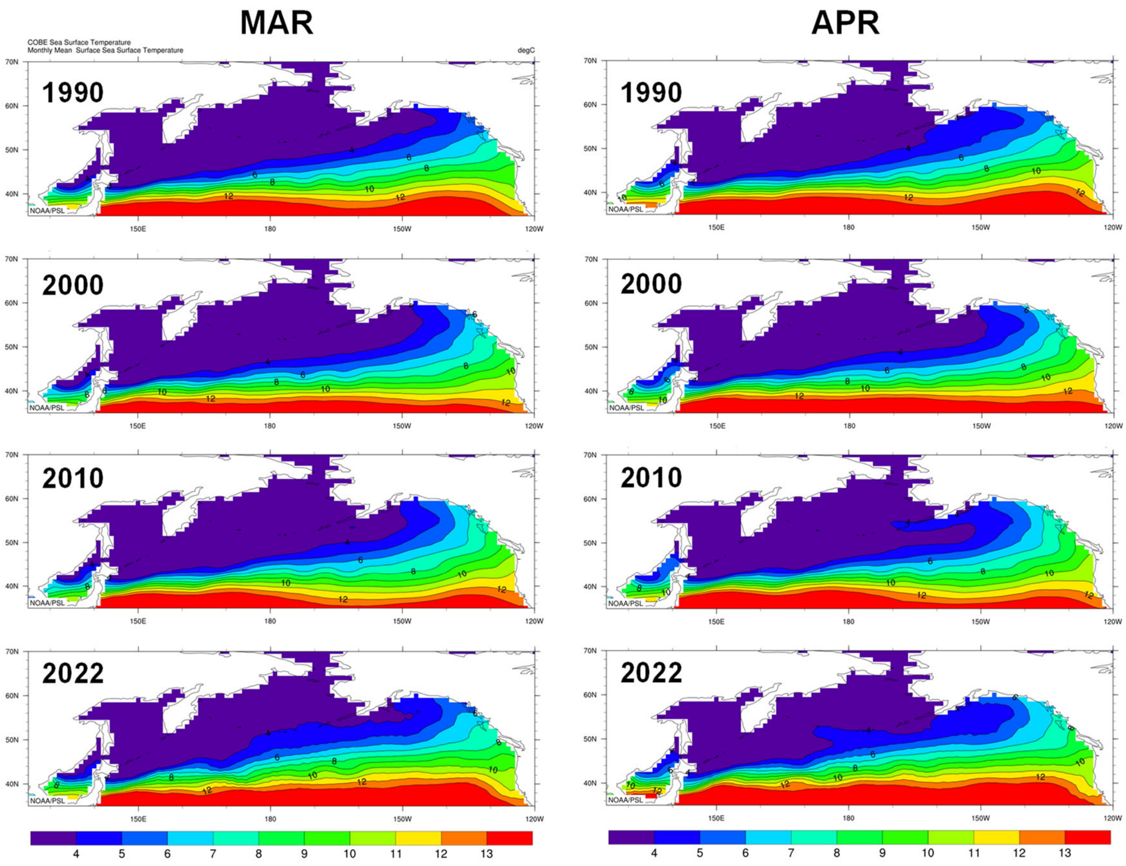


**Figure S5** Northward shift in early spring (March and April) thermal habitat (sea surface temperature isoclines, °C) of chum salmon in the North Pacific Ocean and marginal seas at a decadal scale (1990, 2000, 2010, 2022). Mapped from Plot: COBE Sea Surface Temperature, NOAA Physical Sciences Laboratory. Blue, green, and orange show available temperature range (4–13℃) for chum salmon, based on previous predictions: optimal growth and feeding (8–12℃), available for swimming and feeding (5–13℃), and overwintering temperature range (4–6℃) (Kaeriyama et al., 2012; Urawa et al., 2018). Red (>13℃) and purple (<4℃) indicate potentially critical habitat.

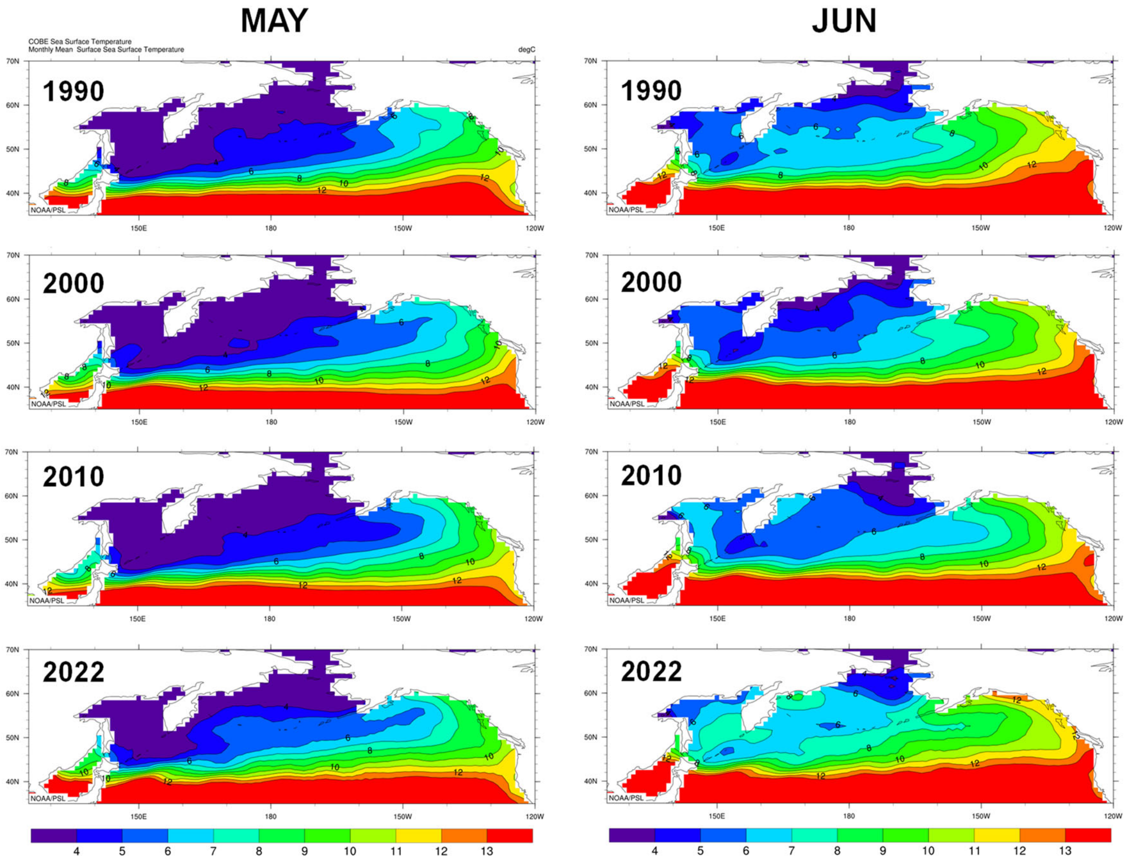


**Figure S6** Northward shift in early summer (May and June) thermal habitat (sea surface temperature isoclines, °C) of chum salmon the North Pacific Ocean and marginal seas at a decadal scale (1990, 2000, 2010, 2022). Mapped from Plot: COBE Sea Surface Temperature, NOAA Physical Sciences Laboratory. Blue, green, and orange show available temperature range (4–13°C) for chum salmon, based on previous predictions: optimal growth and feeding (8–12°C), available for swimming and feeding (5–13°C), and overwintering temperature range (4–6°C) (Kaeriyama et al., 2012; Urawa et al., 2018). Red (>13°C) and purple (<4°C) indicate potentially critical habitat.

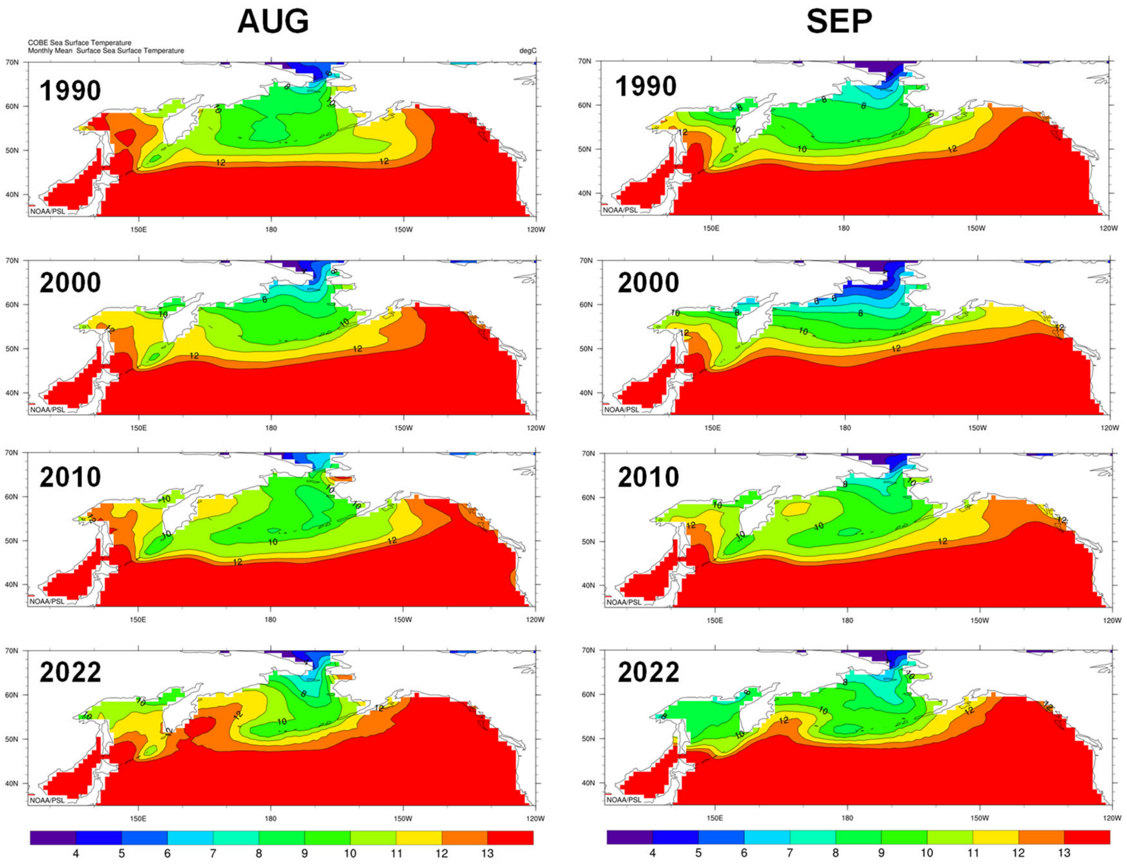


**Figure S7** Longitudinal contraction of high latitude summer (August and September) thermal habitat (sea surface temperature isoclines, °C) of chum salmon in the North Pacific Ocean and marginal seas at a decadal scale (1990, 2000, 2010, 2022). Mapped from Plot: COBE Sea Surface Temperature, NOAA Physical Sciences Laboratory. Blue, green, and orange show available temperature range (4–13°C) for chum salmon, based on previous predictions: optimal growth and feeding (8–12°C), available for swimming and feeding (5–13°C), and overwintering temperature range (4–6°C) (Kaeriyama et al., 2012; Urawa et al., 2018). Red (>13°C) and purple (<4°C) indicate potentially critical habitat.

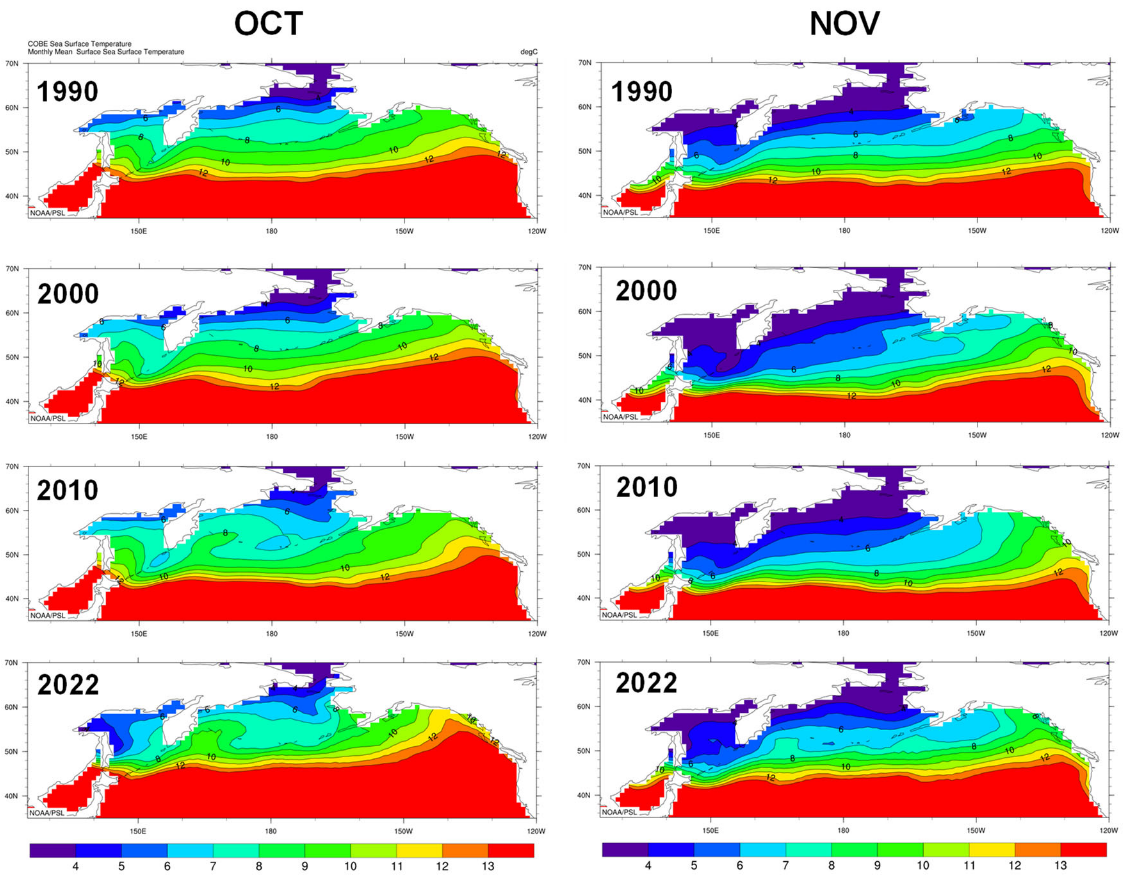


**Figure S8** Northward shift in fall (October – November) thermal habitat (sea surface temperature isoclines, °C) of chum salmon in the North Pacific Ocean and marginal seas at a decadal scale (1990, 2000, 2010, 2022). Mapped from Plot: COBE Sea Surface Temperature, NOAA Physical Sciences Laboratory. Blue, green, and orange show available temperature range (4–13°C) for chum salmon, based on previous predictions: optimal growth and feeding (8–12°C), available for swimming and feeding (5–13°C), and overwintering temperature range (4–6°C) (Kaeriyama et al., 2012; Urawa et al., 2018). Red (>13°C) and purple (<4°C) indicate potentially critical habitat.

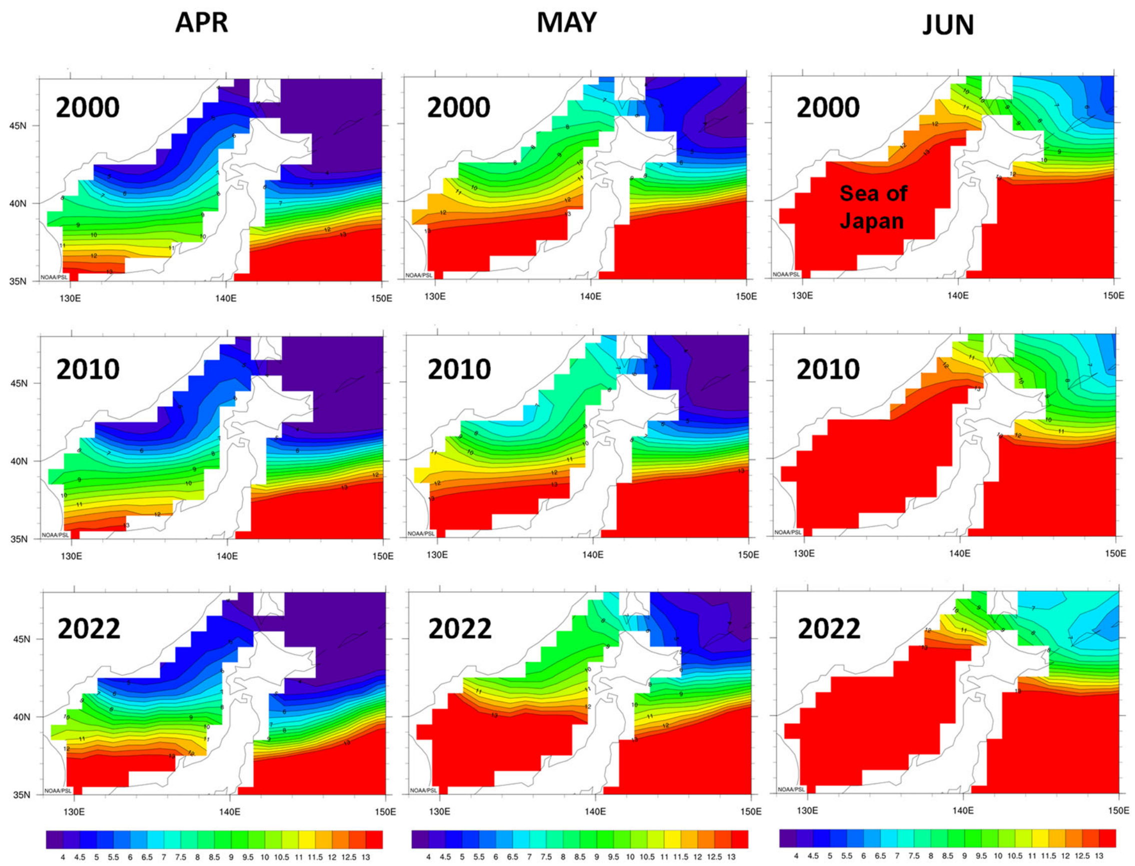


**Figure S9** Northward shift in the spring and early summer (April – June) thermal habitat (sea surface temperature isoclines, °C) of Japanese chum salmon fry in coastal waters surrounding Japan, especially in the Sea of Japan, at a decadal-scale (2000, 2010, 2022). Mapped from Plot: COBE Sea Surface Temperature, NOAA Physical Sciences Laboratory. Blue, green, and orange show available temperature range (4–13℃) for chum salmon, based on previous predictions: optimal growth and feeding (8–12℃), available for swimming and feeding (5–13℃), and overwintering temperature range (4-6℃) (Kaeriyama et al., 2012; Urawa et al., 2018). Red (>13℃) and purple (<4℃) indicate potentially critical habitat.

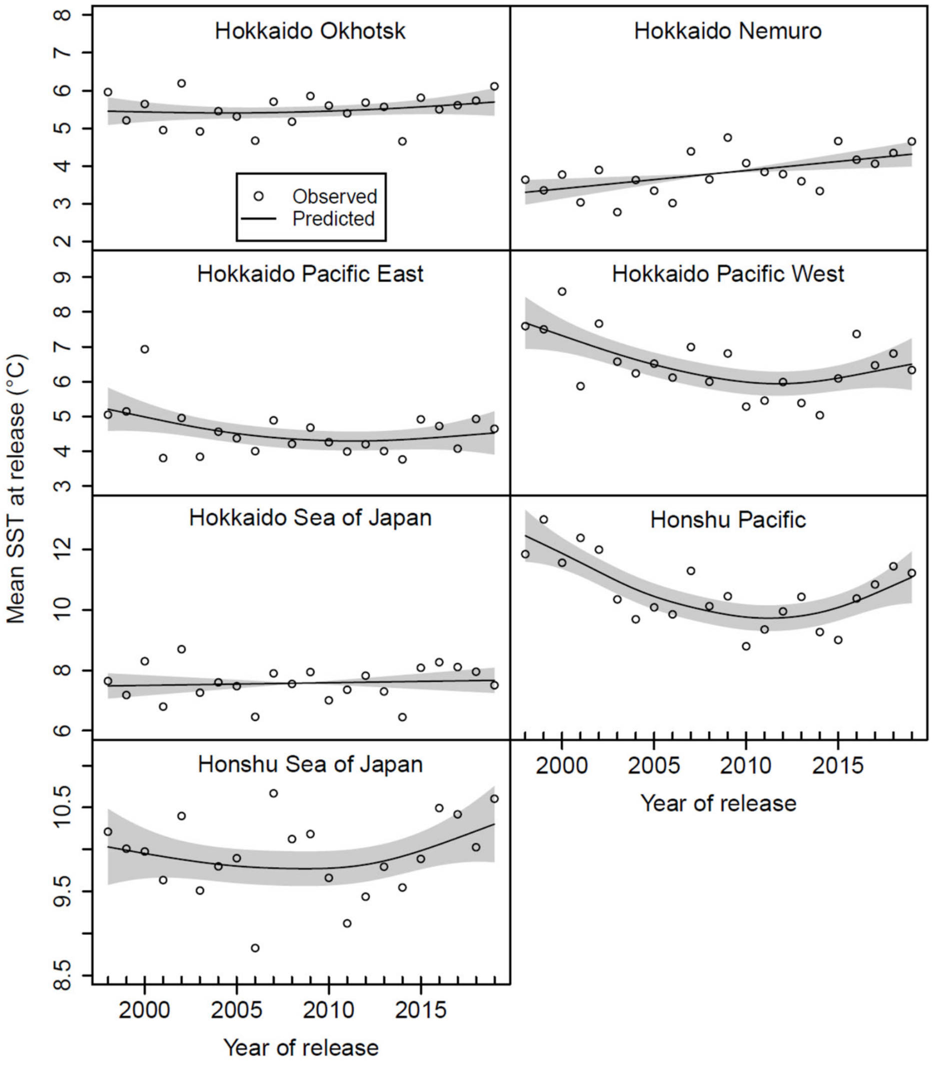


**Figure S10** Annual mean sea surface temperature (SST) at time of release of hatchery chum salmon fry by management region in Japan, 1998–2019, as shown by the nonlinear and/or linear year effect from generalized additive models (GAMs) (Table S1). Shaded areas show 2×standard errors. The annual mean SSTs were calculated by averaging the monthly mean SSTs by sub-region in each management region of Japan: Hokkaido Okhotsk (Abashiri, Monbetsu, and Soya), Hokkaido Nemuro (Nemuro Strait), Hokkaido Pacific East (Nemuro Pacific, Kushiro, and Tokachi), Hokkaido Pacific West (Hidaka, Iiburi, Uchiura Bay, western Tsugaru Strait, and Tsugaru Strait), Hokkaido Sea of Japan (northern and southern Rumoi, Ishikari, western Shiribeshi, and Hiyama), Honshu Sea of Japan (Aomori, northern and southern Iwate, Miyagi, Fukushima, and northern Ibaraki), Honshu Sea of Japan (Aomori, Akita, Yamagata, northern and southern Niigata, and Toyama Bay). See Figure 1a for management region locations.

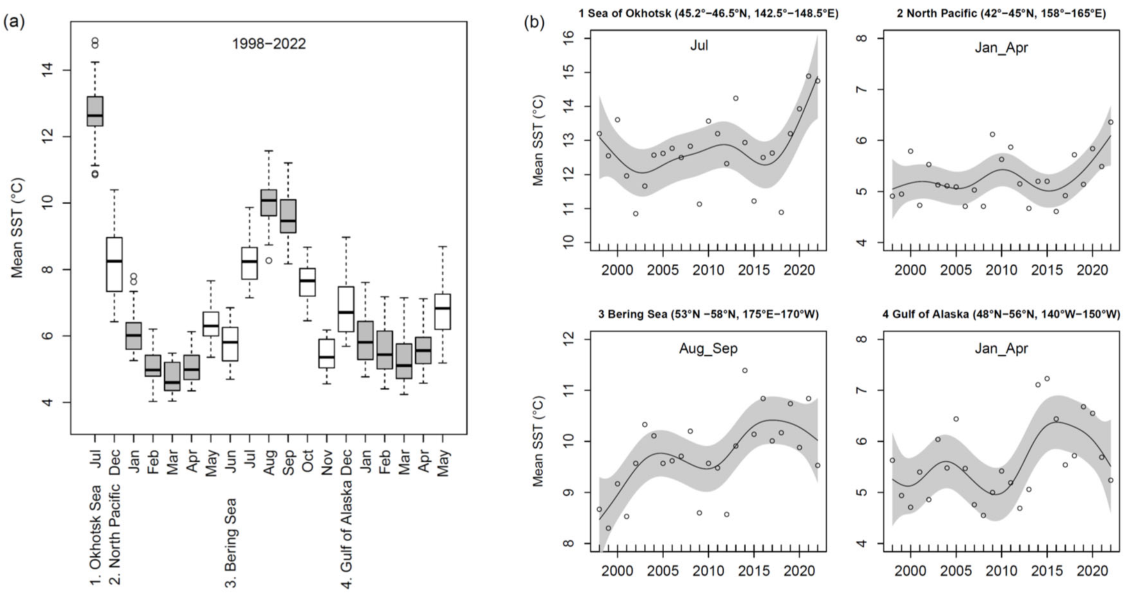


**Figure S11** Mean sea surface temperature (SST) along Japanese chum salmon ocean migration routes (1998–2022); see Fig. 1b, migration areas 1–4). (a) Monthly mean SSTs in feeding and wintering areas in 1998–2022. The gray boxes show the summer and winter SSTs used in our regression model. (b) Summer and winter SSTs used in our regression model. Jan_April and Aug_Sep indicates the mean SSTs during January and April, and August and September, as shown by the nonlinear and linear year effect from generalized additive models (GAMs): Sea of Okhotsk ($p$ = 0.046, $R^2$ = 0.37), western North Pacific ($p$ = 0.118, $R^2$ = 0.29), Bering Sea ($p$ = 0.015, $R^2$ = 0.44), Gulf of Alaska ($p$ = 0.066, $R^2$ = 0.36). Shaded areas show 2×standard errors. The mean SSTs were calculated from the NOAA 0.25° daily Optimum Interpolation Sea Surface Temperature (OISST) climate record.

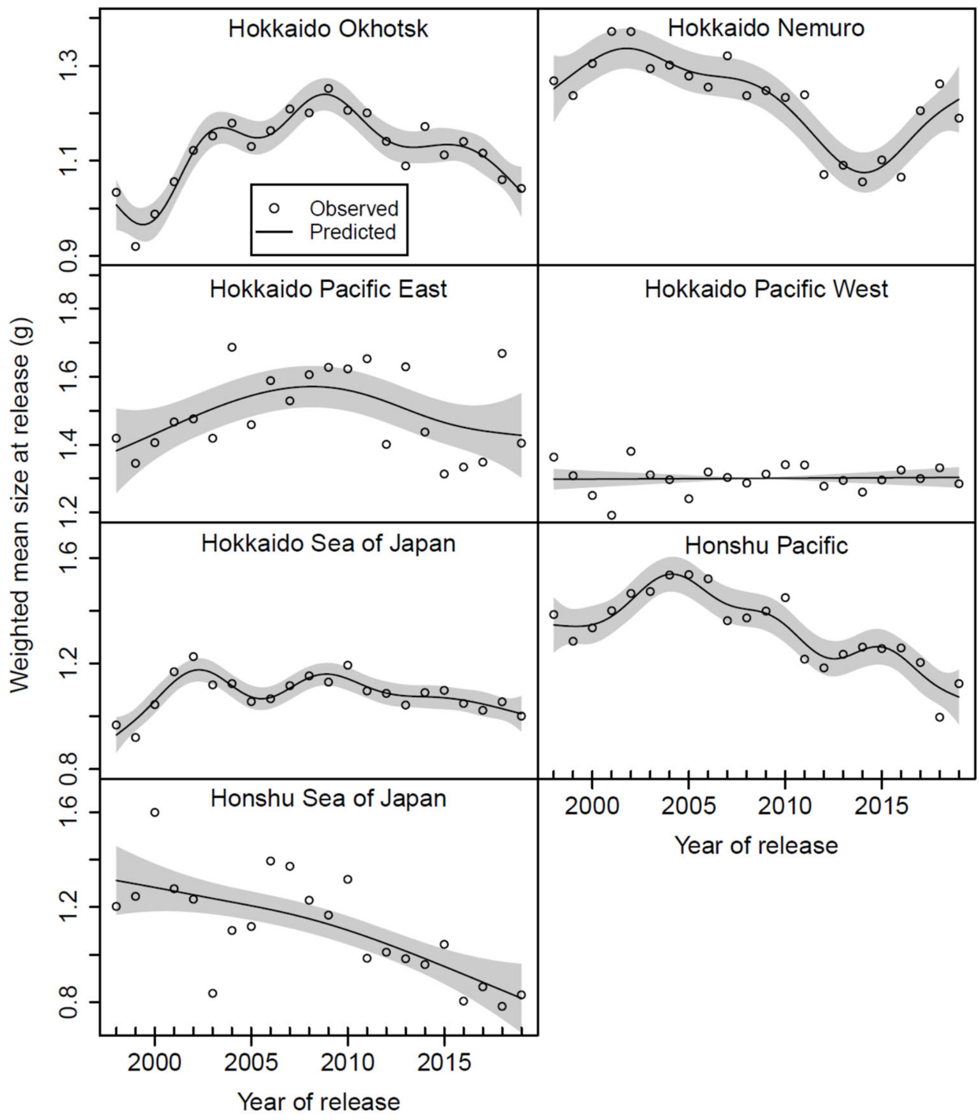


**Figure S12** Mean fry size (body weight in grams) at release of hatchery chum salmon fry by management region in Japan, 1998–2019, as shown by the nonlinear and/or linear year effect from generalized additive models (GAMs) (Table S1). Shaded areas show 2×standard errors. See Figure 1a for management region locations.

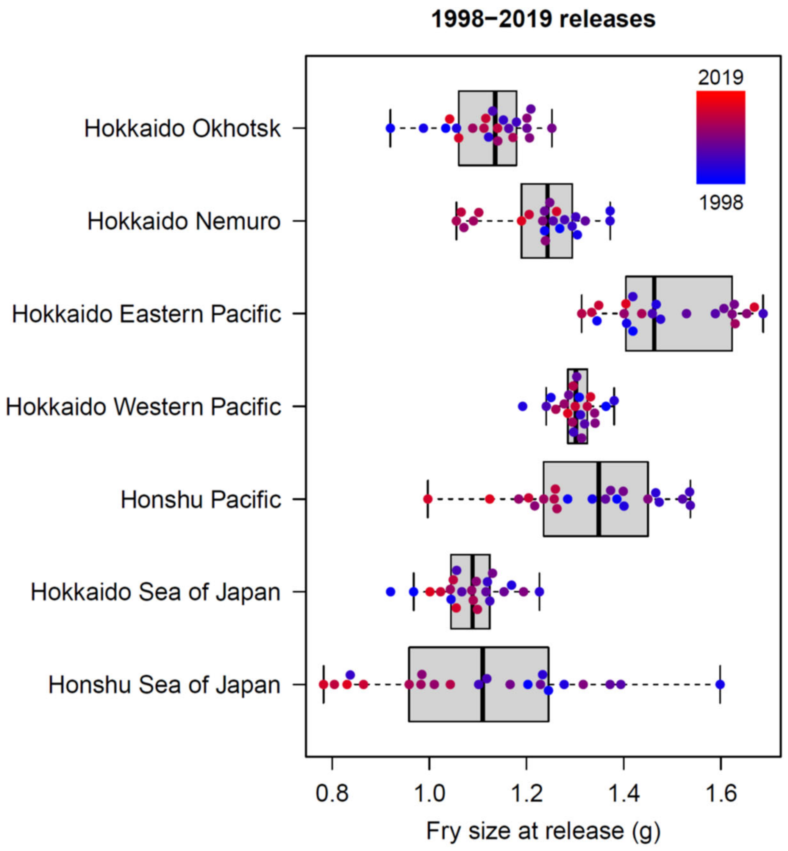


**Figure S13** Fry size (body weight in grams) at release of hatchery chum salmon fry by management region in Japan, 1998–2019. See Figure 1a for management region locations.

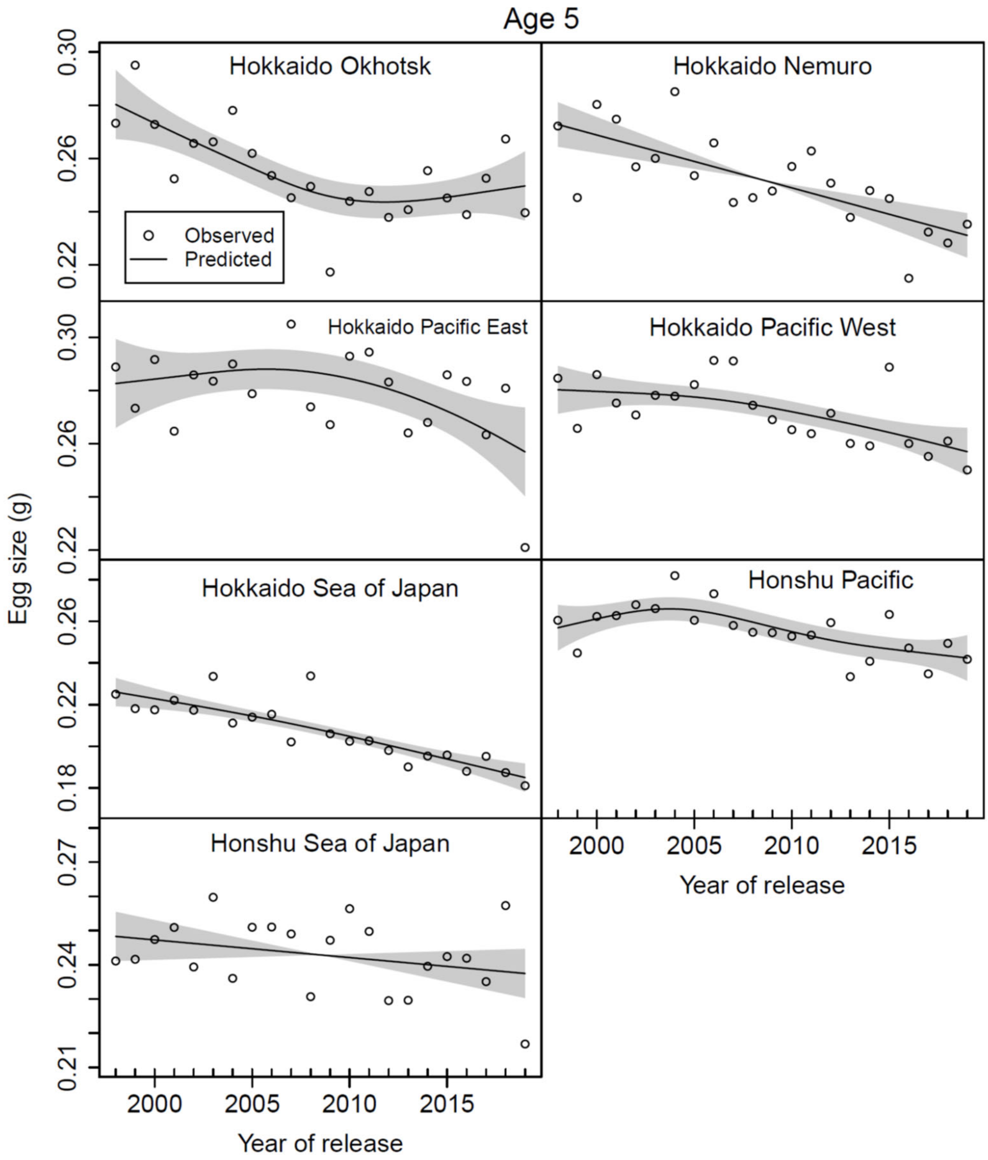


**Figure S14** Mean egg size (weight in grams) of age 5 chum salmon by management region in Japan, used for 1998–2019 releases, as shown by the nonlinear and/or linear year effect from generalized additive models (GAMs) (Table S1). Shaded areas show 2×standard errors. Hokkaido Okhotsk egg size at age 5 was missing in 2005, and we substituted a simple mean of egg sizes in 2004 and 2006. See Figure 1a for management region locations.

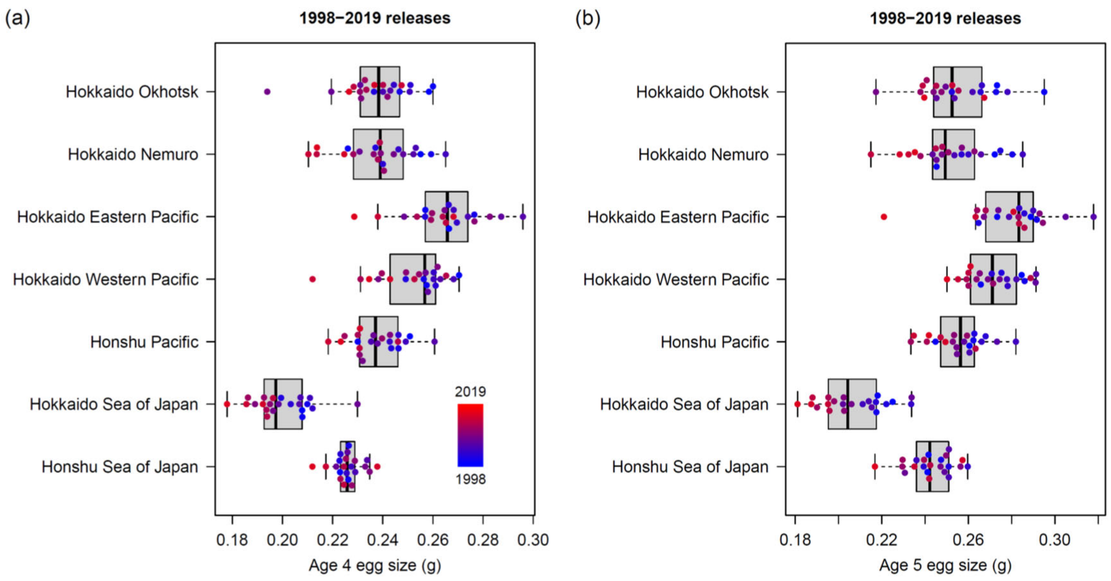


**Figure S15** Mean egg size (weight in grams) of age-4 and age-5 chum salmon by management region in Japan, used for 1998–2019 releases. See Figure 1a for management region locations.

| Explanatory variables | Effect sizes (95% CIs) | Units |
|---|---|---|
| Hokkaido Okhotsk | | |
| 1 Russian pink salmon catch in year t+2 | 0.88 (0.78−1.00) | 100 million fish |
| 2 Yellowtail catch in year t | 0.91 (0.88−0.95) | 100 tons |
| Hokkaido and Honshu Pacific | | |
| 1 Relative fecundity in year t−1 | 0.39 (0.25−0.62) | 100 eggs/FL^3 |
| 2 Russian pink salmon catch in year t+2 | 0.73 (0.62−0.87) | 100 million fish |
| 3 Russian pink salmon catch in year t+3 | 0.80 (0.68−0.94) | 100 million fish |
| 4 Size at release in year t | 1.63 (1.28−2.06) | 0.1g |
| Hokkaido Sea of Japan | | |
| 1 Bering Sea Aug−Sep SST in year t+1 | 0.68 (0.52−0.90) | 1°C |
| 2 Gulf of Alaska Jan−Apr SST in year t+2 | 1.50 (1.17−1.92) | 1°C |
| 3 Okhotsk Jul SST in year t | 0.73 (0.61−0.88) | 1°C |
| 4 Russian chum salmon catch in year t+3 | 0.86 (0.75−0.99) | 10 million fish |
| Honshu Sea of Japan | | |
| 1 North Pacific Jan−Apr SST in year t+1 | 1.30 (1.00−1.68) | 1°C |
| 2 Okhotsk Jul SST in year t | 0.87 (0.77−0.99) | 1°C |
| 3 Russian pink salmon catch in year t+2 | 0.87 (0.76−1.00) | 100 million fish |
| 4 SST at release in year t | 0.62 (0.47−0.81) | 1°C |

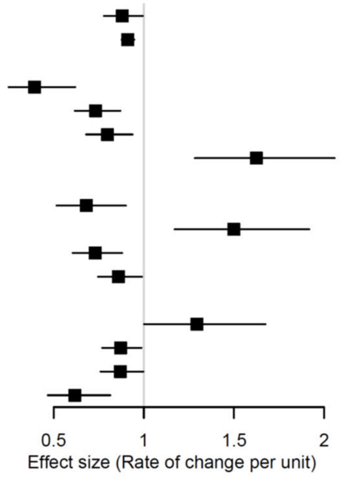


**Figure S16** Practical effect sizes of explanatory variables on Japanese chum salmon marine survival (adult return rates across regions). Effect sizes of best regression models with 95% confidence intervals (exp $(\hat{\beta}_i \pm 1.96 \times \mathrm{SE}(\hat{\beta}_i))$ in four management regions in Japan (Table 3). $t$ = hatchery release year of chum salmon fry. See Figure 1a for management region locations.

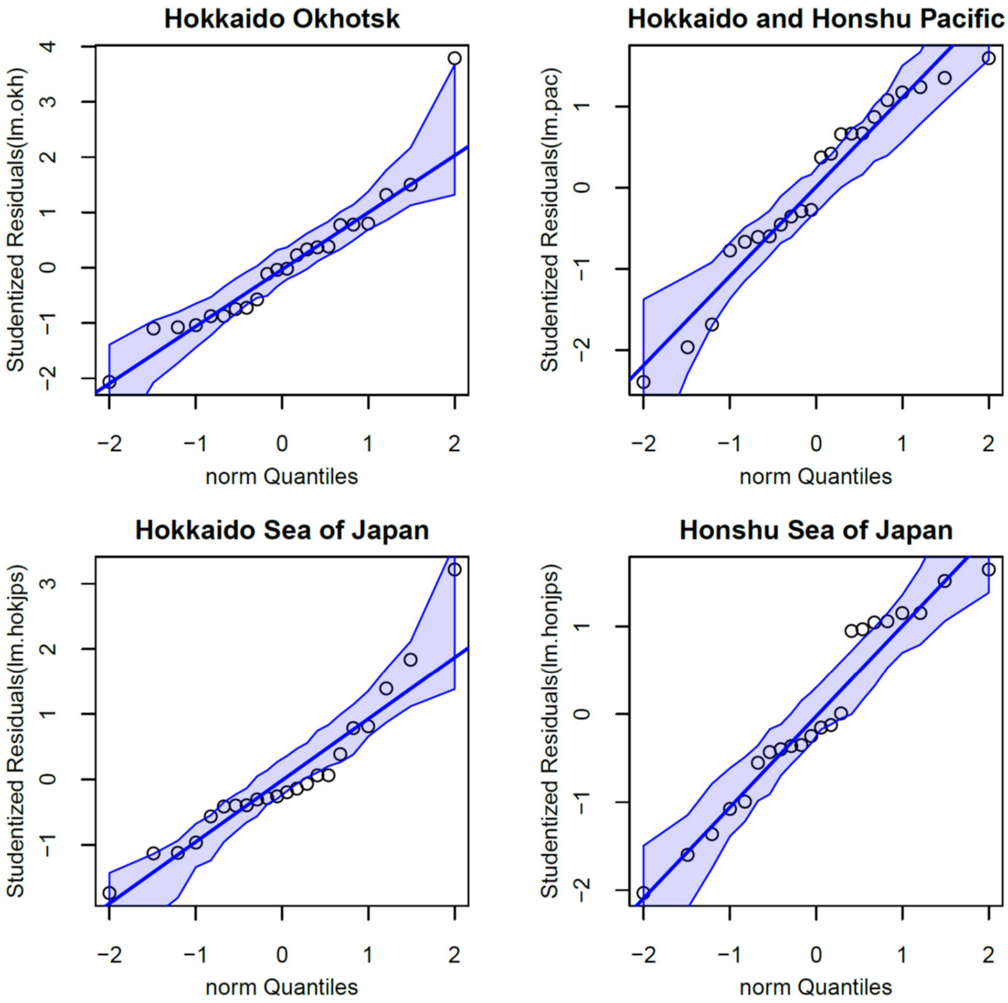


**Figure S17** The qq plots of the residuals from the best models (Table 2) obtained by a normal linear regression of log-transformed return rate using the 'qqPlot' function in the R package car. Shaded areas show pointwise 95% confidence intervals. See Figure 1a for management region locations.

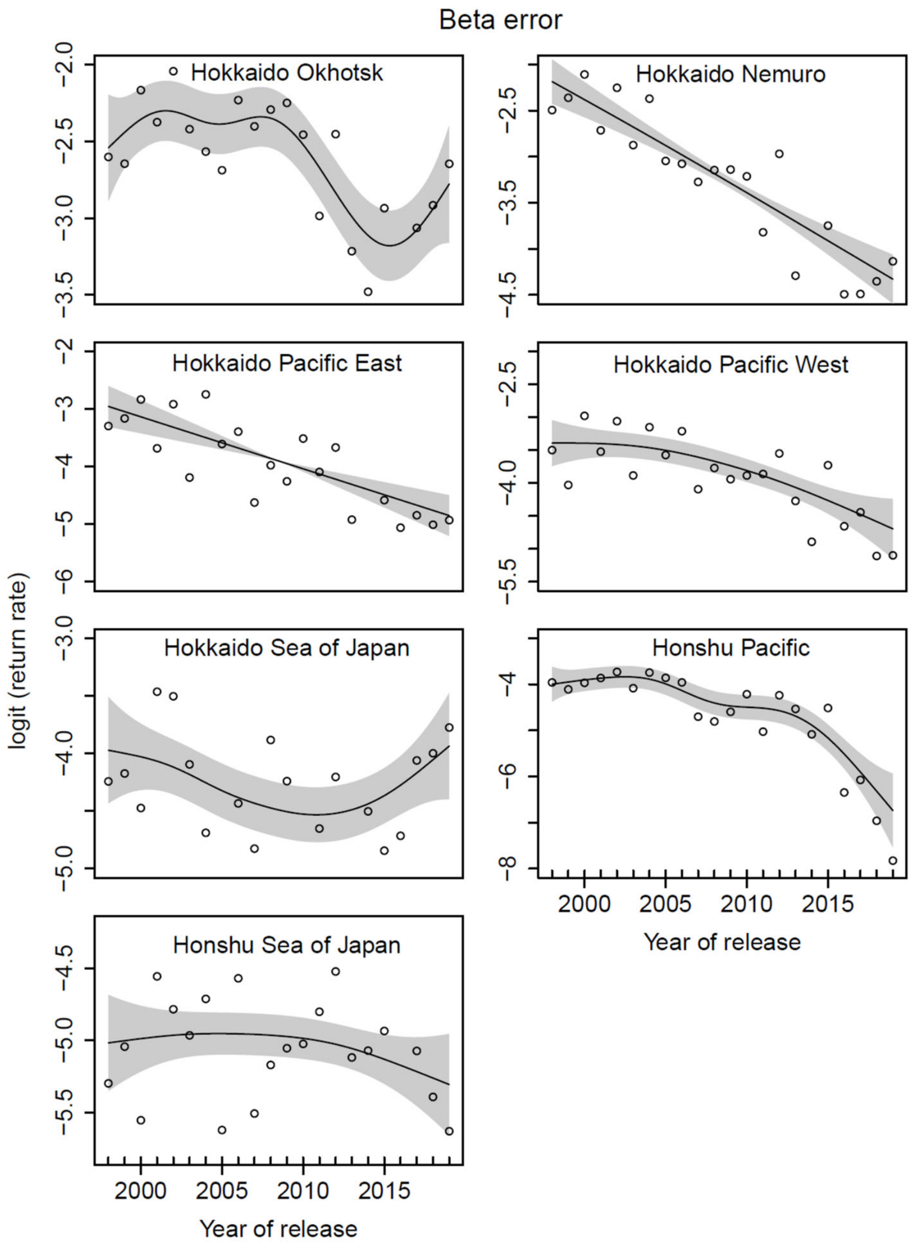


**Figure S18** Logit-transformed chum salmon adult return rates by management region in Japan, 1998–2019, as shown by the nonlinear and/or linear year effect from generalized additive models (GAMs). Shaded areas show 2×standard errors. See Figure 1a for management region locations.

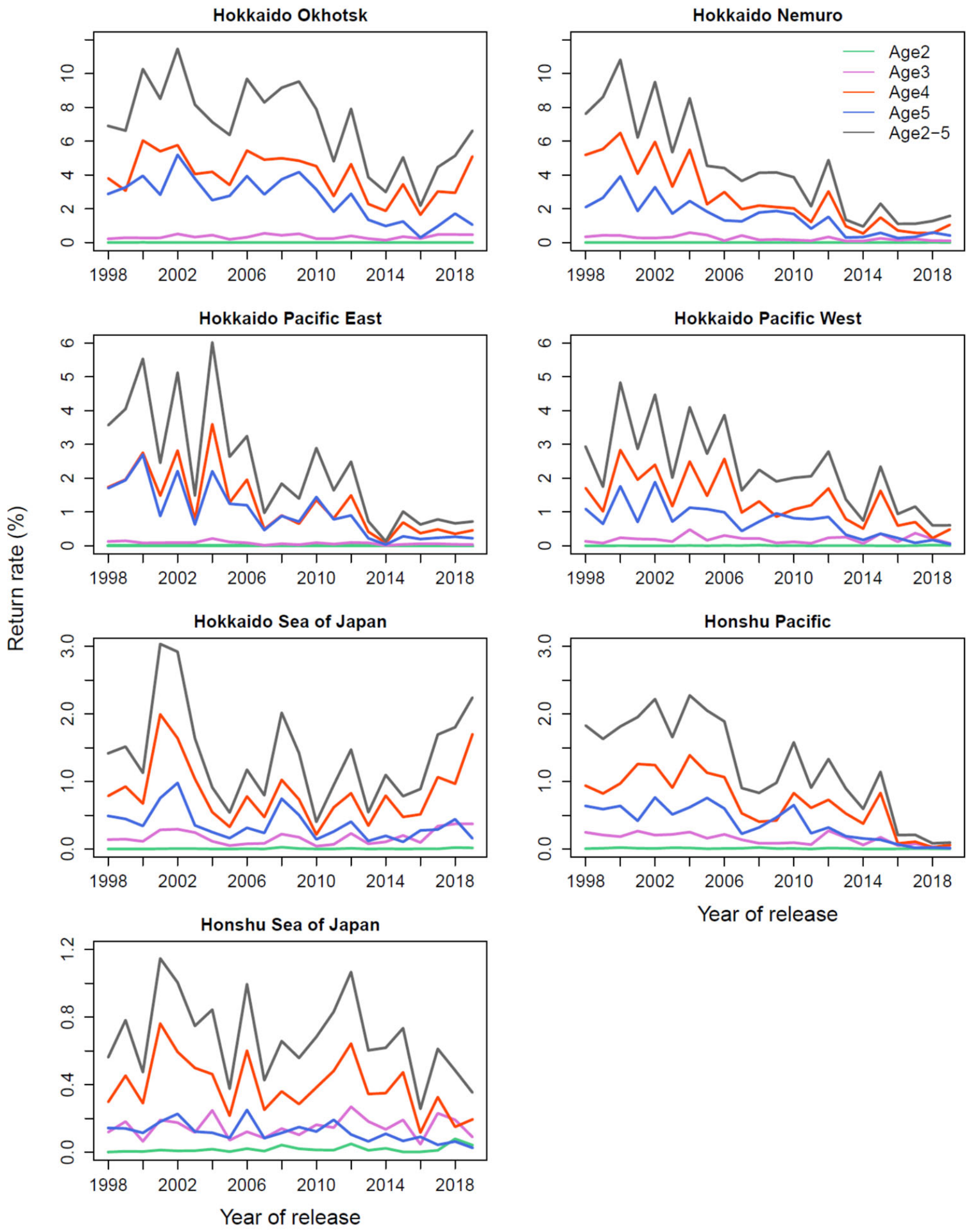


**Figure S19** Age-specific and total adult return rates (marine survival) of chum salmon from 1998–2019 releases in seven management regions in Japan (Figure 1a).

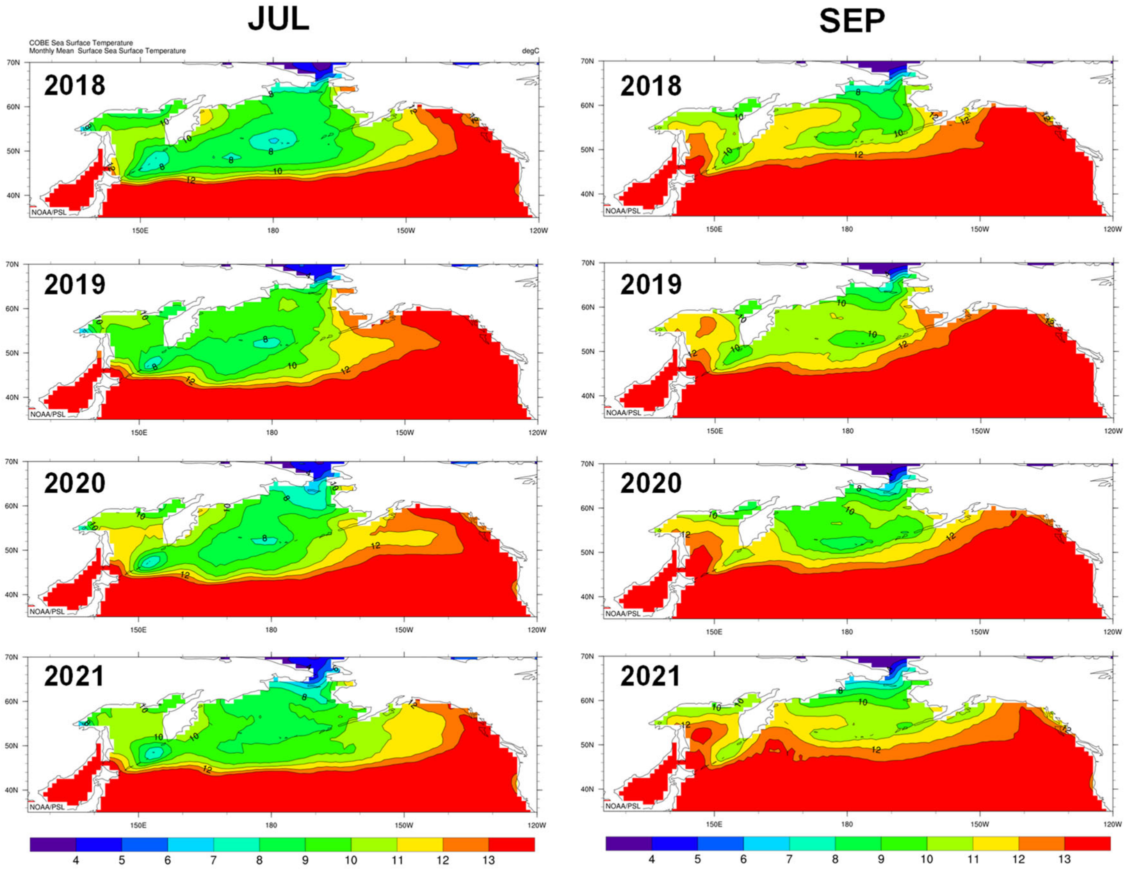


**Figure S20** Annual changes in the July and September sea surface temperature isoclines in the North Pacific Ocean and marginal seas, 2018-2021. Mapped from Plot: COBE Sea Surface Temperature, NOAA Physical Sciences Laboratory. Blue, green, and orange show available temperature range (4–13°C) for chum salmon, based on previous predictions: optimal growth and feeding (8–12°C), available for swimming and feeding (5–13°C), and overwintering temperature range (4–6°C) (Kaeriyama et al. 2012; Urawa et al. 2018). Red (>13°C) and purple (<4°C) indicate critical habitat.

**The R scripts and Data S1–S7**

The following detailed supplementary data are available at https://doi.org/10.5281/zenodo.14868156.

1. The R scripts and data for the regression analysis

2. Supplementary Data S1–S6 in Microsoft Excel files

**Data S1** Size and time of release of Japanese hatchery chum salmon fry in Hokkaido and Honshu (1998–2020), including Hokkaido: 184 rivers, 37,886 release events, $n$ = 21,332,093 fish; Honshu: 141 rivers; 19,605 release events, $n$ = 9,588,413 fish; FRA 2023).

**Data S2** Age composition of adult returns of Japanese chum salmon in Hokkaido and Honshu (1997–2020), including Hokkaido: 52 rivers; Honshu: 102 rivers; 1,902 surveys, $n$=1,210,717 fish; FRA, 2023).

**Data S3** Chum salmon adult female reproductive traits data from 10 rivers in Hokkaido and 14 rivers in Honshu (1997–2020), including egg size (weight), fecundity, fork length, age, and gonad weight ($n$=37,533 fish; FRA, 2023).

**Data S4** Japan coastal 0.25°×0.25° daily mean sea surface temperature (SST) for 18 areas in Hokkaido (1982-2020; JMA, 2022).
https://www.data.jma.go.jp/kaiyou/data/db/kaikyo/series/engan/engandata_SP.html

**Data S5** Japan coastal 0.25°×0.25° daily mean sea surface temperature (SST) for 12 areas in Honshu (1982–2019; JMA, 2022).
https://www.data.jma.go.jp/kaiyou/data/db/kaikyo/series/engan/engandata_SN.html
https://www.data.jma.go.jp/kaiyou/data/db/kaikyo/series/engan/engandata_TK.html

**Data S6** Catch of 56 marine fish species in Japan (1980–2020; MAFF, 2023).

3. Supplementary Data S7 in NetCDF (Network Common Data Form) files

**Data S7** Global 0.25°×0.25° monthly mean sea surface temperature (SST) in four open ocean areas of Japanese chum salmon migration (1982–2022; NOAA, 2023a).
